# High throughput expression-based phenotyping and RNAi screening reveals novel regulators of planarian stem cells

**DOI:** 10.1101/2022.08.29.505550

**Authors:** Erik G. Schad, Christian P. Petersen

## Abstract

The complexity of cell types and states revealed by single-cell RNAseq atlases presents a challenge for the systematic analysis of fate determinants using traditional screening methodologies. Differentiation in the planarian *Schmidtea mediterranea* exemplifies this problem, as these animals continuously produce over 100 differentiated cell types for homeostasis and regeneration using neoblast adult pluripotent stem cells. The signaling factors enabling neoblast self-renewal and selective differentiation of these many fates are still incompletely understood. We developed a method using high-throughput expression profiling by qPCR and whole-animal RNAseq to simultaneously assess numerous cell fate markers as the phenotypic readout in large-scale RNAi screens. Applying this method, we performed an RNAi screen of 400 kinases, receptors, and other regulatory molecules to reveal specific functions for 30 previously unknown factors in neoblast biology. 17 genes were required for neoblast maintenance, including factors likely involved in cell-cycle regulation, nutrient sensing, and chromatin modification. Multidimensional expression information additionally revealed several specific regulators of other neoblast activities, including a *mink1* kinase regulating global neoblast differentiation, the energy responsive kinase *adenylate kinase-2* regulating intestine specification within the neoblast population, an RNA acetyl transferase *nat10* regulating epidermal differentiation, and a *pak1* kinase restricting neoblast localization to prevent tissue outgrowths. These results identify several new regulators of neoblast activities and demonstrate the applicability of expression-based screening for systematic analysis of stem cell phenotypes in whole animals.

## Introduction

Organisms such as planarians, acoels, and hydra that are capable of whole-body regeneration make use of multipotent adult stem cells to generate many or all adult differentiated cells (Srivastava et al., 2014; Tanaka and Reddien, 2011). Understanding how organisms can effectively deploy high potency stem cells in adulthood is therefore an important step in understanding the basis for strong regenerative abilities. In addition, single-cell expression atlases have revealed a large complexity of differentiated adult cell types in animals undergoing whole-body regeneration, raising the question of how such broad cell replacement in adulthood can be accomplished with fidelity (Fincher et al., 2018; Hulett, 2022; Plass et al., 2018; Siebert et al., 2019). High-throughput forward or reverse genetic screens have been valuable in identifying critical regulatory steps in regeneration (Chen et al., 2015; Forsthoefel et al., 2012; Reddien et al., 2005a; Whitehead et al., 2005). However, such approaches are typically limited to examination of overt morphological phenotypes or to a small number of molecular phenotypes because of the limitations of conventional imaging and histology. Given the potential complexity of differentiation pathways available to regeneration, there is a need for methods allowing a higher content readout of cell states that could be implemented at scale in order to efficiently identify control points in stem cell activities for tissue regeneration in vivo.

The process of tissue renewal and regeneration in freshwater planarians uses adult pluripotent stem cells termed neoblasts which contribute to all tissue lineages in adulthood. Neoblasts are present throughout the mesenchyme of the planarian body plan, absent only in the pharynx and head tip (Reddien, 2018). They possess embryonic-like morphology including chromatoid bodies (Coward, 1974), scant cytoplasm, and large nuclei (indicating an undifferentiated state), and are 5-10 microns in size (Reddien and Sanchez Alvarado, 2004). Neoblasts specifically express Piwi proteins (*piwi-1,piwi-2, piwi-3),* other RNA regulatory proteins such as the CELF/Bruno homolog *bruli*, and components of RNA germ granules often associated with germline stem cells in other organisms, including PRMT5, the CCR4-NOT complex, SmB, Tudor, Vasa, and Pumilio factors (Bansal et al., 2017; Fernandez-Taboada et al., 2010; Guo et al., 2006; Kim et al., 2019; Kimoto et al., 2021; Li et al., 2021; Palakodeti et al., 2008; Reddien et al., 2005b; Rouhana et al., 2014; Salvetti et al., 2005; Shibata et al., 2016; Solana et al., 2016; Solana et al., 2009; Wagner et al., 2012). Neoblasts are the only cycling cells in planarians (Morita et al., 1969; Newmark and Sanchez Alvarado, 2000), so they specifically express key factors driving the cell cycle (e.g., *pcna, ribonucleotide reductase)* (Fincher et al., 2018). In addition, their status as the only proliferative cells in planarians enables their purification using FACS with DNA-binding vital dyes such as Hoechst as the cohort of irradiation-sensitive G2/S/M (“X1”) cells (Hayashi et al., 2006; Newmark and Sanchez Alvarado, 2000). Elimination of neoblasts by either RNAi depletion or by ionizing radiation ceases formation of differentiated cells, leading to failed tissue maintenance and ultimately lethality from lysis due to failed epidermal maintenance (Dubois, 1949.; Reddien et al., 2005b; Solana et al., 2012). A combination of irradiation, RNAi, BrdU, transplantation assays, and expression experiments have shown that all differentiated cells in adult planarians are derived from neoblasts (Baguñà et al., 1990; Baguna et al., 1989; Eisenhoffer et al., 2008; Newmark and Sanchez Alvarado, 2000; Reddien et al., 2005b). Transplantation of single neoblasts restore viability and regeneration ability to lethally irradiated animals and contributed to all lineages, demonstrating the pluripotency of cells from this population (Wagner et al., 2011).

Neoblasts are present in several specified states poised for differentiation into distinct lineages (Scimone et al., 2014; van Wolfswinkel et al., 2014). For example, factors such as *zfp-1* and *nkx2.2* are expressed in neoblast subpopulations and these genes are also required for terminal differentiation to epidermis and intestine, respectively (Forsthoefel et al., 2012; van Wolfswinkel et al., 2014; Wagner et al., 2011). In many cases, such specification factors are expressed both in neoblast subpopulations as well as in terminally differentiated cells, for example the *ovo* transcription factor expressed in both neoblasts and mature eyes and required for eye differentiation (Lapan and Reddien, 2012). By contrast, in some lineages, post-mitotic progenitor cells with unique transcriptional states have been identified as likely cellular intermediates undergoing further differentiation (Eisenhoffer et al., 2008). For example, epidermis formation likely involves specification of *zfp-1+* neoblasts, followed by exit from proliferation and formation of *prog1+* progenitor cells, which then mature further through *AGAT+* progenitors and ultimately terminally differentiate into mature epidermal cells (Cheng et al., 2018; Tu et al., 2015; van Wolfswinkel et al., 2014). Transitions between neoblast specification states are also likely possible. The isolation and transplantation of a neoblast subpopulation of *tetraspanin+* cells can repopulate irradiated animals with descendant cells that restore the complement of neoblast subtypes and also restore regeneration and homeostasis ability (Zeng et al., 2018). In addition, the analysis of neoblast composition in 2-cell clones revealed that neoblasts randomly adopt new specified states during each new S phase, suggesting neoblast pluripotency may arise from an intrinsic fate-switching mechanism linked to cell cycle transit (Raz et al., 2021). Regeneration involves the use of a similar set of neoblast-specified subpopulations present prior to wounding (Benham-Pyle et al., 2021), indicating a close association between the differentiation programs used by neoblasts for regeneration and homeostatic renewal.

The processes governing neoblast abilities are still incompletely understood, but regulatory factors of controlling neoblasts have been identified that control aspects of self-renewal, differentiation, and migratory targeting. Piwi factors facilitate differentiation by surveilling transposons activated during the lineage-specific activation of large transcriptional regions of the genome (Li et al., 2021). Neoblast activities are also controlled by chromatin-modifying factors such as CHD4 and the NuRD complex, MLL factors controlling early and later differentiation and PRC2 controlling self-renewal (Dattani et al., 2018; Duncan et al., 2015; Hubert et al., 2013; Jaber-Hijazi et al., 2013; Mihaylova et al., 2018; Scimone et al., 2010; Vasquez-Doorman and Petersen, 2016; Wagner et al., 2012). The CELF protein *bruli* controls neoblast self-renewal and an alternative splicing program in neoblasts (Guo et al., 2006; Solana et al., 2016). Assays to measure neoblast repopulation from clones following sublethal irradiation determined roles for transcription factors *soxP-1* and *junli-*1, and RNA-binding factors *KHD-1* and *CIP-29*, in self-renewal (Wagner et al., 2012). Regulation of neoblasts involves canonical cell cycle factors such as *repA1, ANAPC1*, and *Rb* (Pearson and Sanchez Alvarado, 2010; Reddien et al., 2005a) and input into proliferation/differentiation outputs through other regulatory factors such as PTEN, Akt, and TOR (Oviedo et al., 2008; Peiris et al., 2016; Peiris et al., 2012; Tu et al., 2012). A *mex3*-dependent regulatory step enables neoblasts to exit from proliferation in order to differentiate down multiple lineages (Zhu et al., 2015). Neoblasts undergo migration to target locations in the animal, induced by injury and also in order to maintain particular differentiated tissues in the absence of injury (Guedelhoefer and Sanchez Alvarado, 2012). This process is dependent on integrins and also the EMT-regulating transcription factors *snail-1/-2* and *zeb-1* (Abnave et al., 2017; Bonar and Petersen, 2017; Seebeck et al., 2017) and is coupled to DNA damage repair processes (Sahu et al., 2021). Niche factors for neoblasts are not yet fully elucidated, but based on the phenotypes from disruption of intestine formation, this organ may be a source of signals important for neoblast maintenance (Forsthoefel et al., 2012). Planarian muscle also constitutes a major source of ECM that provides additional signals for controlling neoblast activity (Chan et al., 2021; Cote et al., 2019; Lindsay-Mosher et al., 2020). Furthermore, regional signals can guide neoblast differentiation, for example an anterior Wnt/Notum regulatory loop regulates neural progenitors (Hill and Petersen, 2015), Hedgehog signaling guides neural specification (Currie et al., 2016), EGF signals specify differentiation of intestine, eyes and other tissues (Barberan and Cebria, 2019; Barberan et al., 2016; Emili et al., 2019; Fraguas et al., 2011), and BMP specifies epidermal progenitors into dorsal versus ventral identity (Wurtzel et al., 2017). *egfr-3* and its putative ligand *nrg-7* are required for self-renewal during neoblast repopulation after sublethal irradiation (Lei et al., 2016). However, neoblasts are not eliminated in homeostatic *egfr-3(RNAi)* animals, suggesting additional factors may act in parallel to fulfill the total mitogenic signals necessary for normal neoblast maintenance. Together, a variety of signals influence neoblast activity, but the core regulation dictating proliferation and differentiation decisions is not yet fully understood.

To seek out additional factors required for planarian stem cell function, we developed an alternative approach by conducting a large-scale RNAi screen modified to use expression profiling by qPCR and RNAseq as a phenotypic readout after inhibition of candidate regulatory factors. The methodology enabled assessment of key stem cell and progenitor states after inhibition, defining functions for genes in the differentiation pathway, and identifying new control points for tissue homeostasis in planarians.

## Results

### Development of a high-throughput RNA extraction method for large-scale gene expression analysis in planarians

We devised a high-throughput RNAi screen to probe the roles of 398 kinases and other signaling molecules in neoblast-dependent tissue renewal (Figure 1A). dsRNAs targeting each gene were administered by feeding 6 times over 3 weeks, following transverse amputation and scoring for regeneration defects 8 days later (Figure 1B, Figure S1, Table S1). Regeneration defects were qualitatively scored by blastema extent and shape as described previously (Reddien et al., 2005a) and binned into overall categories of “death”, “no regeneration”, “reduced regeneration”, and “unaffected.” dsRNA targeting the *Photinus pyralis* luciferase gene (not present in the planarian genome) resulted in normal regeneration (Figure 2B). Several dsRNA positive control conditions were included in the screen, including *smad4* and *bruli*, which resulted in expected reduced blastema phenotypes (Figure 1A-B) (Guo et al., 2006; Reddien et al., 2005a). Overall, 23.1% of RNAi treatments caused regeneration defective phenotypes, with 5.5% resulting in death, 11.3% with severe regeneration defects, and 6.3% with weak regeneration defects (Table S1), similar to frequencies from prior screens (Reddien et al., 2005a). Genes recovered in the screen were involved in a variety of cellular processes, including cell proliferation (*Rb*, *cdk2, cdk12,* etc.), transcriptional regulation (*mnat, cdk7),* RNA splicing and posttranscriptional regulation (*prpf4b, nat10*), metabolic regulation (*ak2, papss2, pank3*), and signal transduction (e.g., *pkn2, mink1, pak1, pipp5k2*, *lamtor, akt*) (Figure 1B). Taken together, this screen identified multiple previously unknown factors required for proper regeneration in planarians.

**Figure 1.**
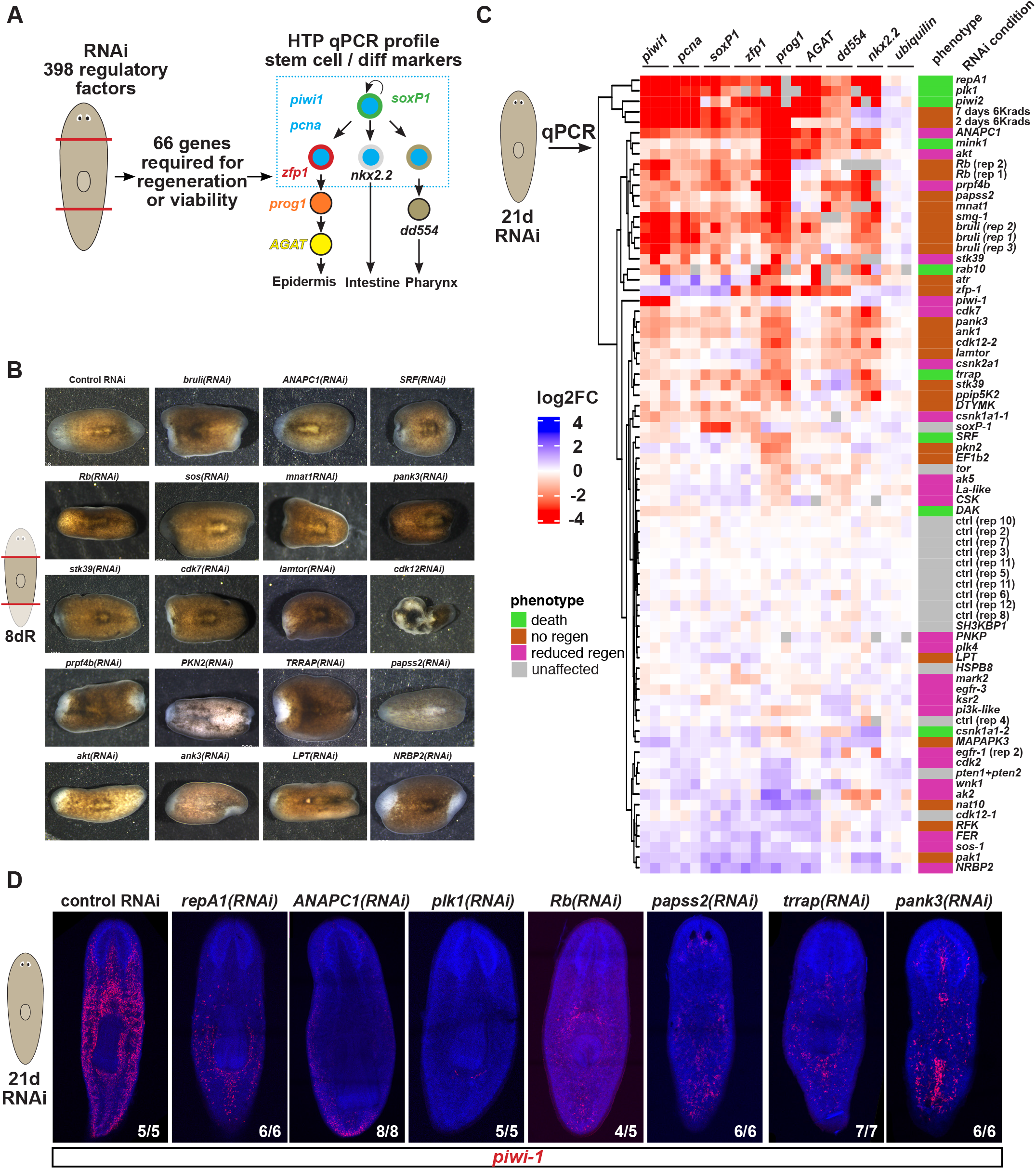
Combining high-throughput expression profiling and RNAi screening to find regulators of planarian stem cells. (A) Screen design showing inhibition of 398 kinases and other regulatory factors to identify 66 required for regeneration and viability. Animals were dosed with dsRNA by 6 feedings over 2 weeks followed by amputation and regeneration for 8 days prior to morphological scoring. Genes required for regeneration or viability were re-screened through homeostatic RNAi followed by collection of whole-animal RNA in biological triplicates and quantification by high-throughput qPCR of expression of mRNAs marking key neoblasts (*piwi-1, pcna, soxP-1*), and fate markers for several planarian cell lineages (epidermal: *zfp-1, prog-1, agat-1*; intestine: *nkx2.2*, pharynx: *dd554*). Right diagram denotes abbreviated planarian cell lineage with relevant markers indicated. (B) Examples of live phenotypes of regeneration deficiency isolated from primary RNAi screen representing majority phenotype from 10 animals tested in each condition. Blastema tissue is initially unpigmented, allowing assessment of blastema size, morphology, and presence of newly formed eyes. (C) qPCR expression profile from secondary screen in which animals were inhibited for each gene through 6 dsRNA feedings over 21 days. Rows indicate RNAi conditions or control treatments with lethal doses of ionizing irradiation (6,000 Rads) known to deplete neoblasts and RNA collected 2 or 7 days later as indicated. Columns indicate markers probed by qPCR with the biological replicates shown. Heatmap shows log2 fold-change expression values normalized to the average control dsRNA conditions within several experimental batches (each labeled as “ctrl”). Expression values were determined by the delta-delta-Ct method using *clathrin* as a housekeeping normalizing control. Expression of a second negative control marker *ubiquilin* shows the specificity of treatments on neoblast/progenitor expression states. Gray boxes indicate no detection by qPCR. Log2 fold-change values, standard deviations, p-values and adjusted p-values are shown in Table S2. Right heatmap shows regeneration phenotype severity scored from the primary screen. Biological replicate conditions for *bruli* RNAi*, Rb* RNAi, and control (“ctrl”) RNAi are denoted with replicate number (eg, rep 1, rep 2, etc). Genes not previously defined in planarians are named after the best blastx match to the human proteome. (D) FISH of *piwi-1* in uninjured animals following RNAi of select genes recovered in the screen (6 dsRNA feedings over 21 days). Numbers in panels indicate animals with expression as shown.

**Figure 2.**
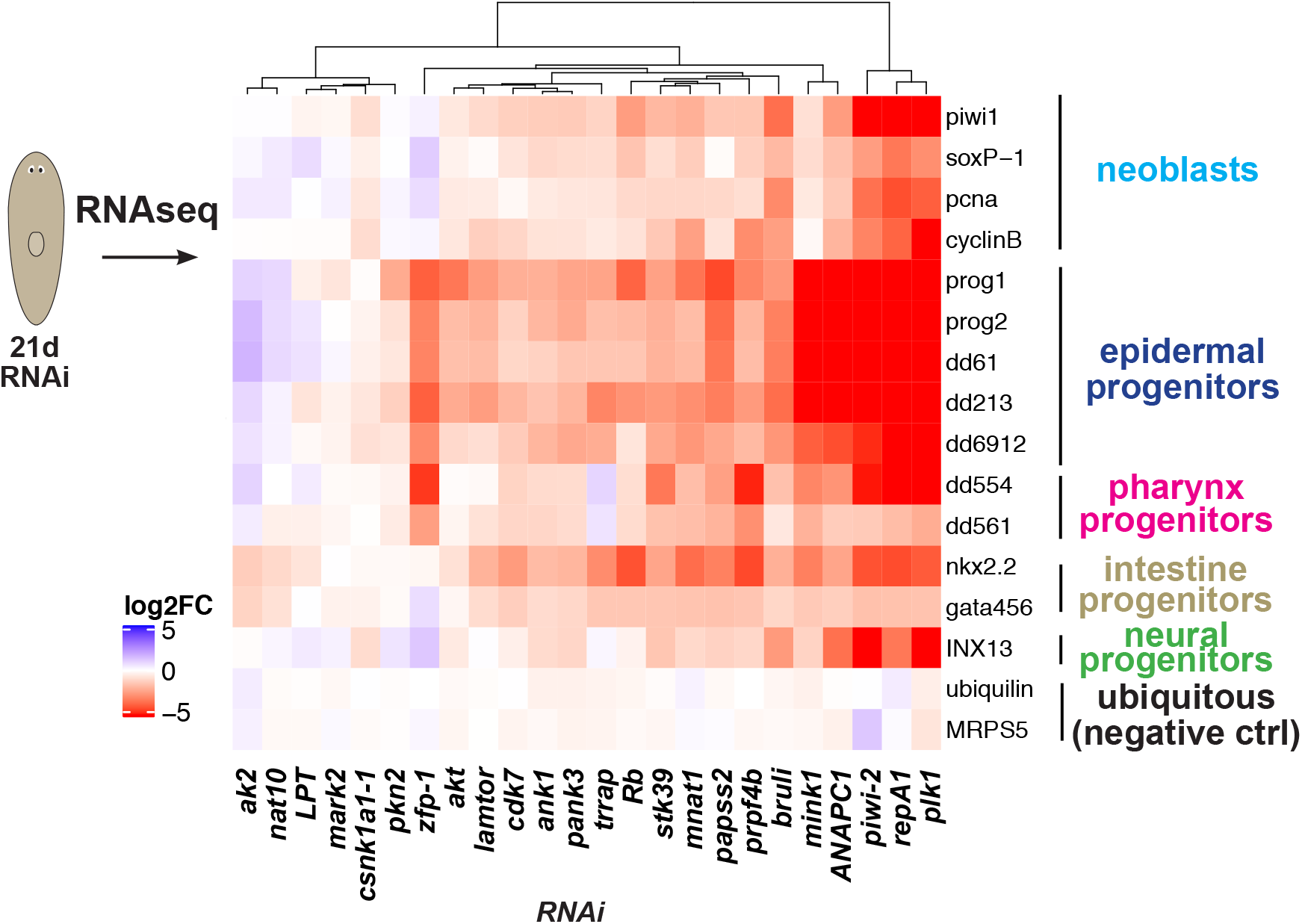
High-throughput RNAseq to define stem cell phenotypes in an RNAi screen for regeneration deficiency. RNAseq was performed on 25 of the conditions used in the secondary screen with three biological replicates for each condition and normalized to control RNAi conditions. Heatmap shows average log2 fold-change from three biological replicates for several neoblast and progenitor markers, compared to negative Control RNAi conditions targeting the *Photinus pyralis* luciferase gene not present in the planarian genome. *ubiquilin* (dd1364) and *mitochondrial ribosomal protein S5* (MRPS, dd1692) are ubiquitously expressed genes not predicted to be altered by RNAi of tested genes. Columns show RNAi treatments, rows show marker gene expression. Right labels show category of marker for reporting Full dataset expression log2 fold-change expression values, and adjusted p-values shown in Table S3.

In addition to scoring visible phenotypes following RNAi, we sought a methodology to examine the expression of multiple factors marking both neoblasts and several types of differentiating post-mitotic progenitor cells. Fluorescence in situ hybridizations or other histological analyses have been used in whole-animal functional screens to identify factors responsible for the normal formation or maintenance of particular cell types in planarians. However, these methods are not readily amenable to detection of more than 2-3 mRNA markers simultaneously and do not provide quantitative information about expression levels. We took a different approach and developed a high-throughput RNA extraction and qPCR assay (Figure 1A, S2A). After RNA extraction, expression status of any key cell state marker could in principle be ascertained to guide discovery of factors required for specific aspects of tissue renewal. We adapted an RNA extraction protocol using proteinase-K digestion for homogenization (Ly et al., 2015), which allows rapid homogenization of planarian tissue without requiring a tissue grinder. Trizol extractions were then performed in 96-well plates, and RNA purified using 96-well nucleic acid binding columns (Figure S2A). cDNA generated from this RNA was suitable for detection of key neoblast and progenitor genes uniquely marking key transcriptional states in differentiation of abundant differentiated cell types that we reasoned would be easily detected in this manner: *piwi-1* and *pcna* marking neoblasts, *soxP-1* marking a large neoblast subpopulation*, zfp-1* marking a neoblast subpopulation believed to be specified for epidermal fate*, prog-1* marking early post-mitotic epidermal progenitors*, agat-1* marking later post-mitotic epidermal progenitors believed to be matured from *prog-1+* cells*, dd554* marking post-mitotic progenitors of the pharynx*, nkx2.2* marking both gamma neoblasts specifying toward intestine as well as mature intestine cells, and *clathrin* as a normalizing control. Omission of the reverse transcriptase step resulted in substantially less qPCR signal, indicating successful detection of mRNA for each probe (Figure S2B).

As an additional test of the utility of these probes for monitoring neoblast-dependent cell states, we verified that expression of these markers was reduced by 6000 Rads of ionizing radiation, known to eliminate neoblasts within 1-2 days, followed by successive depletion of any post-mitotic progenitors having rapid turnover. We tested a dose series of irradiation from 850 to 6000 Rads then harvested RNA for detection of neoblast transcripts by qPCR. All irradiation dosages reduced neoblast and differentiation markers, including *piwi-1* (also known as *smedwi-1*, a pan-neoblast marker), *soxP-1* (sigma marker), *zfp-1* (zeta-marker), *prog-1* (post-mitotic epidermal progenitors), and *nanos* (labeling the nascent testes germline stem cells present but not able to mature in asexual planarians) (Figure S3A). Stronger doses such as lethal irradiation (6000 rads) caused correspondingly stronger reduction to the expression of these markers. Next, we tested RNAi controls for expected gene expression changes. *bruli* inhibition resulted in reduction to all neoblast markers and also differentiating progeny, *smedwi-1* inhibition only reduced *smedwi-1* expression, *soxP-1* inhibition reduced *soxP-1* expression, and *zfp-1* inhibition reduced *zfp-1* and developmentally downstream epidermal differentiation marker *prog-1* expression. As a negative control, inhibition of *tolloid* did not affect any marker expression (Figure S3B). These results verified the expression detection methodology for use in subsequent analysis of planarian whole-animal RNAi phenotypes.

We selected a subset of 60 of 398 genes from the primary screen that were required for optimal regeneration for further analysis through a secondary screen to analyze the expression of neoblasts and markers of epidermis, intestine, and pharynx differentiation by qPCR. We found in pilot experiments that challenging neoblast-deficient animals to regenerate often resulted in their lethality, either preventing analysis of their RNA or requiring gene-specific selections of time-points for RNA collection. To circumvent this issue, we performed the secondary screen under homeostatic RNAi in the absence of injury, testing factors whose inhibition caused lethality or regeneration defects in the primary screen. We performed RNAi using 6 dsRNA feedings and a waiting period for a total of 21 days, and subsequently extracted RNA and performed qPCR in high throughput (Figure 1C). Any RNAi condition that had resulted in death in the primary screen was monitored and RNA collected along with RNAi negative controls at the time when early phenotypes of lethality presented themselves (for example, head regression, lesions, or ventral curling). After cDNA synthesis, we performed high-throughput qPCR on the following neoblast markers in each RNAi condition: *smedwi-1, pcna* (pan-neoblasts), *soxP-1* (sigma neoblasts), *zfp-1* (zeta-neoblasts), *prog-1, agat-1* (epidermal differentiation markers), *nkx2.2* (gamma-neoblasts and lowly expressed in mature intestine cells), *clathrin* (as a normalizing control), and *ubiquilin* (housekeeping marker that should not change after alteration to neoblasts) (Figure 1C).

Hierarchical clustering of the dataset revealed a complex set of expression states generated by inhibition of genes from the screen (Figure 1C, Table S2). As an internal validation of the screen approach, RNAi conditions targeting *piwi-1, soxP-1*, and *zfp-1* were included, and each treatment reduced the corresponding transcripts. In addition, *bruli* dsRNA was included in each of 3 experimental batches encompassing the secondary screen, and *Rb* dsRNA was included in two different batches. In confirmation of expectations from prior work, each treatment caused robust neoblast marker downregulation, with replicate conditions each placed adjacently through hierarchical clustering all RNAi treatments in the screen. Together, these observations suggest the fidelity of the qPCR approach in assessing gene expression changes for critical stem cell and progenitor markers in planarian RNAi screens.

The screen revealed several categories of defects to self-renewal and differentiation following RNAi. A large category of gene inhibitions caused reduction of *piwi-1* neoblast marker expression, including *repA1*, *plk1*, *piwi-2*, *ANAPC1, smg-1, prpf4b, Rb, papss2, pank3, ank1,* and *csnk1a1-1* (at least 2-fold downregulated at padj<0.1 corrected for false discovery by the Benjamini-Hochberg method) (Figure 1C, Table S2). In addition, RNAi of other factors caused downregulation of *piwi-1* to a lesser extent (less than 2-fold downregulated, padj<0.1), including *cdk12-2*, *lamtor*, *csnk2a1*, *trrap*, *SRF*, *ak5* and *dak* (Figure 1C, Table S2). RNAi of other factors (*mnat1*, *stk39*, *rab10*) caused on average strong (>2-fold) downregulation of *piwi-1+* transcripts but were not statistically significant after false-discovery correction (padj>0.1), which could either be a result of false discovery, of higher real variability of *piwi-1* expression across biological replicates due to incomplete penetrance, or of higher detection variability from the qPCR assay format. The dataset also allowed a systematic comparison between severity of regeneration/viability defects and effects on homeostatic levels of *piwi-1* neoblast marker expression. Many phenotypes of regeneration dysfunction and lethality occurred with reductions to *piwi-1* expression, but some strong regeneration defects did not (eg., *ak2*, *pak1*, *LPT*) (Figure 1C, Figure S4). By contrast, treatments resulting in neoblast or progenitor upregulation were rare and had lower effect sizes, indicating that positive regulators of neoblast expression are in general much more abundant than negative regulators (Figure 1C, Table S2).

The average log2 fold-change expression changes of neoblast markers *pcna* and *soxP-1* closely resembled those of *piwi-1* (Figure 1C). Therefore, genes required for normal expression levels of these markers are likely involved in neoblast maintenance through self-renewal rather than controlling the gene expression of individual neoblast markers. Some genes in this category are associated with cell cycle control (*plk1, cdk12, repA1, ANAPC1*), expected because neoblasts are the only cycling cells in planarians and require proliferation for renewal. We also found a variety of other functions required for stem cell maintenance. *ankyrin-1* (*ank1*) encodes an integral membrane protein underlying the spectrin-actin cytoskeleton and may regulate cell motility, proliferation, and cell contacts (Cunha and Mohler, 2011), all of which are likely important for neoblasts. *3’-phosphoadenosine 5’-phosphosulfate synthase 2* (*papss2*) encodes a kinase important for the regulation of sulfation, mediating a number of processes including Snail-mediated cell migration (Xu et al., 2001). Finally, *pantothenate kinase 3* (*pank3*) initiates the first step of coenzyme A biosynthesis, suggesting energy status may play a role in planarian stem cell self-renewal (Ni et al., 2002). As predicted from the known relationship between neoblasts as the source of progenitor cell types across lineages, in the majority of conditions in the screen, downregulation of neoblasts also corresponded to downregulation of markers for early epidermis (*prog-1*), pharynx (*dd554*), and/or intestine lineages (*nkx2.2*) (Figure 1C, Table S2).

Treatments causing reduction of progenitor markers without affecting neoblasts (progenitor marker >2-fold downregulated and padj < 0.1; *piwi-1* expression padj > 0.1) were rare and included RNAi of *zfp-1*, *mink1*, *akt*, *pkn2*, *ak2,* and *mark2*. *zfp-1* is known to be required for epidermal differentiation, and our analysis found *zfp-1* RNAi caused mild upregulation to other neoblast markers (*piwi-1*, *pcna*, *soxP-1*) while decreasing epidermal progenitors *prog-1, agat-1* but not intestinal progenitor marker *nkx2.2*, consistent with prior results. *zfp-1* RNAi also downregulated *dd554* pharynx progenitor marker, suggesting this gene promotes formation of both epidermis and at least some cell types of the pharynx. *mink1* RNAi caused significant (padj<0.1) and much stronger downregulation of all progenitor markers *prog-1*, *AGAT-1*, *dd554* and *nkx2.2* than of neoblast markers and is a candidate for a regulator of differentiation across diverse lineages. *akt* and *pkn2* RNAi reduced epidermal marker abundance without affecting intestine or pharynx progenitors, indicating they are particularly important for epidermal differentiation. *pkn2* encodes a protein kinase known to interact with AKT signaling, and regulates cell-cycle progression, cytoskeleton remodeling, and cell migration (Bourguignon et al., 2003; Lee et al., 2016). By contrast, inhibition of *ak2* selectively reduced *nkx2.2* intestine progenitor expression, suggesting a specific role in differentiation of that lineage. Furthermore, inhibition of *mark2*, a member of the Par-1 family of cell polarity proteins, specifically reduced expression of *zfp-1*. The lack of downregulation of post-mitotic epidermal progenitor markers in this condition could reflect an early stage in the phenotypic progression after inhibition of this gene, or alternatively a separate role for *zfp-1* in whole animals.

Other phenotypic classes were recovered but were rare. Two treatments resulted in downregulation of neoblast markers without substantial downregulation of progenitor markers. For example, RNAi of a *dak,* a homolog of dihydroxyacetone kinase 2 acting in glycerol catabolism, caused weak (less than 2-fold) but significant downregulation of *piwi-1* and *pcna* without affecting other markers. Inhibition of a casein kinase homolog, *csnka1-1,* significantly downregulated *piwi-1* and *zfp-1* expression by ∼2-fold and downregulated *pcna* and *soxP-1* more weakly (<2-fold) but did not have strong effects on progenitor expression. Factors in this category could have been sampled at early phenotypic stages of impaired self-renewal prior to effects on progenitors, or alternatively, these conditions could modify rates of differentiation in conjunction with neoblast abundance. Another class of defects recovered in the screen included phenotypes of progenitor upregulation, which were also rare. A cluster of three RNAi treatments (*cdk2*, *wnk1*, *pten1/pten2* double RNAi) caused a similar phenotype of weak *pcna* and *prog-1* upregulation (less than 2-fold upregulated, padj<0.1). RNAi of planarian PTEN homologs was previously found to cause hyperproliferation, and our results suggest *cdk2* and *wnk1* may similarly control neoblast cell cycle progression and thereby selectively impact expression of these two markers. Finally, a small number of RNAi conditions led to increased expression of neoblast markers. Only one of these, *pak1* RNAi, caused increased expression to more than one neoblast marker. In addition, the effect sizes of treatments causing neoblast marker downregulation were much larger than those causing neoblast marker upregulation. Together, the methodology revealed several classes of phenotypes for further investigation.

We used data from a planarian cell atlas to cross-reference expression patterns of the cell type markers and also the genes identified in the screen (Fincher et al., 2018) (Figure S5A-B). Some genes from the screen, primarily those known to be involved in cell cycle control, had expression highly enriched in neoblasts (e.g., *repA1*, *plk1*, *ANAPC1*, *Rb*). Others showed partial enrichment in neoblasts and also other cell populations (e.g., *atr*, *dak*, *pkn2*, *pknp, mnat1, papss2*), and most other factors were expressed broadly across several cell populations including in neoblasts. Therefore, the action of such factors on neoblast renewal or differentiation could either be cell autonomous or non-autonomous via other contributing tissues. Interestingly, the planarian scRNAseq cell atlas indicates *pank3* is strongly and specifically expressed in intestine and Cathepsin+ cells and only very weakly in neoblasts, suggesting its function in Coenzyme A biosynthesis from these tissues could perhaps nonautonomously contribute to the microenvironment necessary for neoblast maintenance.

To begin to validate the effects observed by qPCR analysis, we first used in situ hybridizations to verify that inhibition of seven genes caused reduced expression of *piwi-1* (Figure 1D). Inhibition of positive control genes *repA1*, *ANAPC1,* and *Rb* caused downregulation of *piwi-1* predicted from their known requirements for regeneration and neoblast maintenance, as well as their known involvement in cell cycle progression. Inhibition of *plk1*, *papss2*, *trrap*, and *pank3* recovered from the screen also caused reductions to neoblast abundance as measured by *piwi-1* staining.

To further validate the conclusions made by qPCR from the screen, we subjected 25 conditions (3 biological replicates each) to expression analysis by RNA-seq including control-treated dsRNA, using preparation of libraries in 96-well format. An advantage of using high-throughput RNAseq as a readout for RNAi screens is that in principle many additional markers could be simultaneously assessed, and the data can be reanalyzed to draw additional conclusions as new markers of cell types/states are uncovered. We first examined several markers of neoblasts, epidermis, pharynx, and intestine recovered from prior functional studies and the planarian cell atlases, which included all of the markers tested by qPCR (Figure 2, Table S3). The RNAseq profiles broadly matched those measured by qPCR, with several conditions resulting in downregulation to neoblast markers (eg, *plk1, repA1, piwi-2, papss2, prpf4b, bruli, Rb, pank3, ank1, cdk7, lamtor, csnk1a-1*), those required only or more strongly for epidermal progenitors (*zfp-1*, *pkn2, akt*), and those required for multiple types of progenitor expression but not strongly for neoblast expression (*mink1*). We next surveyed the expression behavior of a larger number of markers of cell types defined by the planarian cell atlas (Fincher et al., 2018)(Figure S6, Table S3 and Table S4). This analysis confirmed that phenotypes of neoblast downregulation broadly also corresponded to phenotypes of downregulation of markers for several progenitor classes (epidermis, intestine, pharynx). Other planarian lineages such as mature neurons and parenchymal cells likely undergo slower turnover and/or lack progenitor cell types that can be uniquely marked by individual transcripts, and therefore did not substantially downregulate during the limited time frame of animal survival following stem cell depletion or dysfunction. Taken together, our qPCR and RNAseq analysis identify broad regulators of neoblast maintenance, as well factors required more specifically in specification and differentiation.

### *mink1* kinase is required for neoblast differentiation down multiple lineages

We focused subsequent analysis on a more detailed study of factors from the screen displaying unique phenotypic characteristics. A category of genes from the screen caused strong reductions to markers of various differentiating progenitors without as extensive changes to markers of neoblasts after their inhibition, including *misshapen-like kinase 1 (mink1), protein kinase N2 (pkn2)*, and *akt.* Inhibition of these three genes strongly reduced progenitor expression while neoblast markers remained unchanged suggesting they specifically promote differentiation of neoblasts. We first confirmed by FISH that *pkn2* RNAi caused downregulation of *prog-1* expression without affecting neoblast expression (Figure S7). We next examined *mink1* because this factor appeared required for multiple types of progenitor cell markers.

*mink1* encodes a serine/threonine kinase possessing an S/T kinase domain (evalue = 1.9E-86, SMART domain prediction) as well as a CNH domain (evalue = 4.62E-47) common to NIK-family kinases that include Drosophila *misshapen*, *C. elegans mig-18*, and mammalian MINK1, TNIK1, and NIK germinal center kinase family members. NIK kinases have roles in activation of the JNK and p38 pathways, cytokinesis, and non-canonical Wnt signaling (Daulat et al., 2012; Hu et al., 2004; Hyodo et al., 2012). *mink1(RNAi)* animals ultimately underwent lysis and death, so we inhibited the gene for a shorter timeframe to analyze the phenotypic progression of loss of differentiation. 3 dsRNA feedings over 2 weeks of RNAi resulted in lesions and ultimately death, a phenotype previously linked to lack of production of mature epidermis and subsequently loss of epidermal integrity (Figure 3A). We analyzed expression of several cell type markers by qPCR following *mink1* inhibition and found that neoblast factors (*smedwi-1, soxP-1,* and *zfp-1)* were only minimally affected (Figure 3B). However, markers of epidermal differentiation including *prog-1* and *agat-1* were very strongly decreased. Additionally, markers of progenitor cells from the intestine (*nkx2.2*), pharynx (dd554), and markers of germline stem cells (*nanos*) were also strongly decreased following *mink1* RNAi. Using FISH, we confirmed that in *mink1(RNAi)* animals, *smedwi-1+* cells were only slightly reduced in abundance, while *prog-1* cells were almost completely eliminated in uninjured animals (Figure 3C). This suggests *mink1* is likely broadly needed for differentiation down many or most cell lineages. We examined additional requirements for *mink1* in differentiation across multiple lineages as predicted by the qPCR and RNAseq data. First, we examined *ovo,* a transcription factor required for the production of photoreceptor cells that is expressed in both mature eyes and also migratory eye progenitors honing to their destination in two anterior trails (Lapan and Reddien, 2012). Inhibition of *mink1* eliminated the *ovo+* trails of photoreceptor progenitor cells, but not preexisting mature photoreceptor cells (Figure 3D). In concordance with *mink1’s* requirement for *nkx2.2+* intestine progenitor expression, we found that *mink1(RNAi)* animals had reduced staining of *porcupine+* differentiated intestine cells (Forsthoefel et al., 2012). Finally, we examined a marker of pharynx progenitor cells (*dd554*), which was also reduced in number following inhibition of *mink1(RNAi)* (Figure 3D). Taken together, these results suggest that *mink1* exerts a broad control over differentiation down multiple lineages descended from planarian neoblasts.

**Figure 3.**
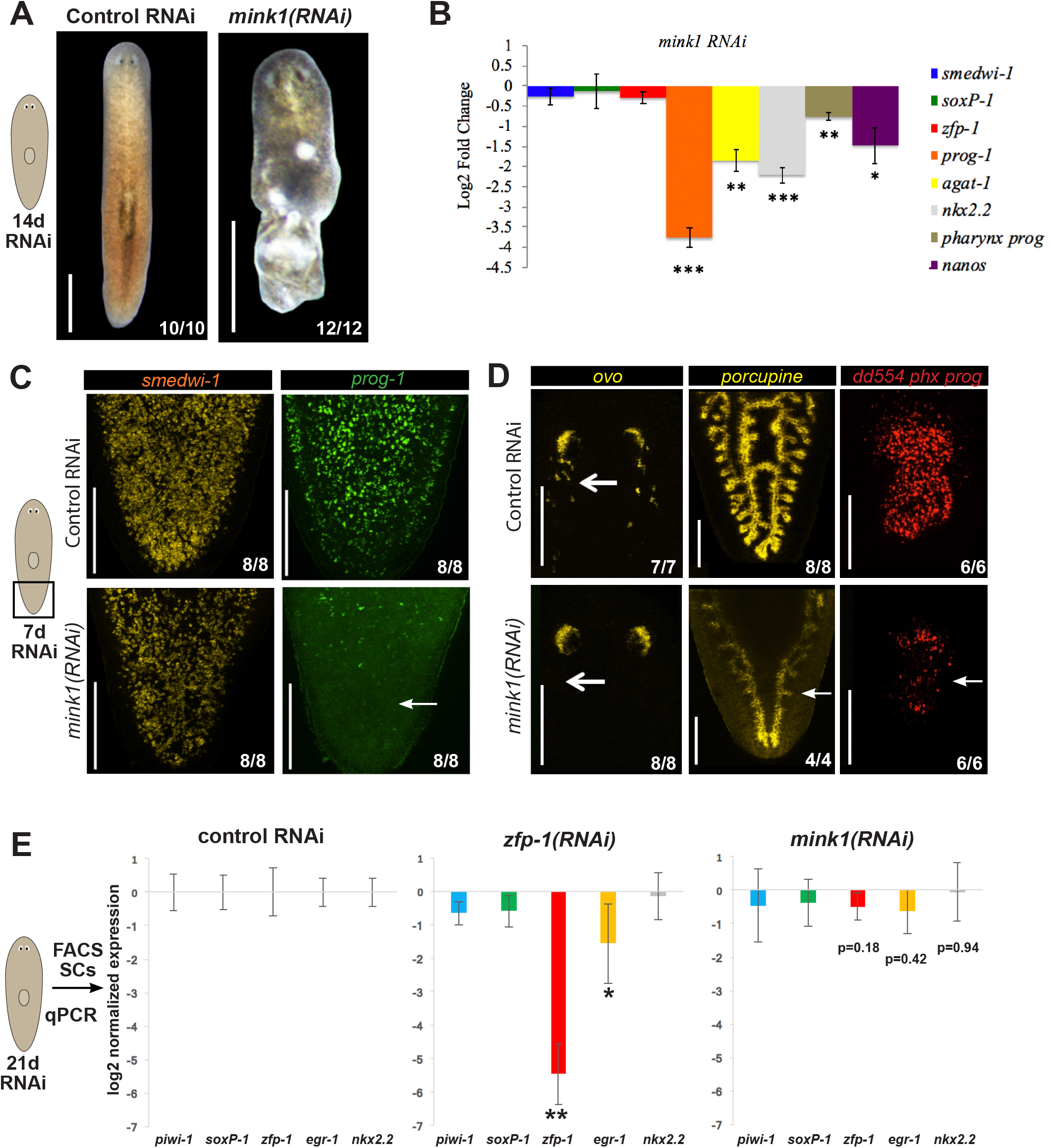
*mink1* kinase is required for production of multiple post-mitotic progenitor cell types. (A) *mink1* RNAi caused lesions in uninjured animals preceding animal lysis and death. (B) Average log2 fold-change qPCR expression in whole-animal *mink1(RNAi)* versus control dsRNA from primary screen, plotted from data in Figure 1C and Figure S2. Bars show averages of 3 biological replicates normalized to control dsRNA conditions, and error bars show standard deviations. ** p<0.01; ***p<0.001 by unpaired two-tailed t-tests. *mink1* RNAi caused strong downregulation of progenitor markers from epidermis (*prog-1, agat-1)*, intestine (*nkx2.2*), pharynx (*dd554*), and germline cells (*nanos*), but not markers of neoblasts. (C) Animals homeostatically inhibited for *mink1* versus control dsRNA and stained by FISH for *piwi-1* (*smedwi-1*) and *prog-1* expression (arrows). *prog* expression was eliminated without loss of *piwi-1* expression in 8/8 animals across two replicate experiments. (D) *ovo* expression marking trail of eye progenitor cells (top arrow), *porcupine* marking differentiated intestine cells, and *dd554* marking pharynx progenitor cells. *mink1* RNAi caused reduction to *prog-1*, migrating *ovo+* cells, reduction to mature gut marker expression, and reduction to pharynx marker expression (arrows). (E) Planarian G2/S/M cells of the “X1” population were FACS sorted in biological replicate from control RNAi, *zfp-1(RNAi)*, and *mink1(RNAi)* animals and markers of neoblast subpopulations analyzed by qPCR. Bars represent averages of 3 biological replicates, 4 animals per sample, bars represent standard deviations, * represents p < 0.05 and ** p < 0.01 from unpaired two-tailed t-tests. *zfp-1* RNAi caused downregulation of *zfp-1* and *egr-1* transcripts marking epidermal specifying neoblasts. By contrast, *mink1* RNAi did not affect these or expression of intestine-specifying neoblasts expressing *nkx2.2*. Bars, 800 (A) or 300 microns (C-D).

The whole-animal qPCR and RNAseq data suggested that *mink1* likely does not act at the level of ensuring specification states within the neoblast population, because *mink1* inhibition did not strongly alter levels of *zfp-1* but in contrast reduced expression of post-mitotic markers such as *prog-1*. As further confirmation of this mechanism, we examined expression of specified neoblast states using qPCR from FACS-sorted G2/S/M neoblasts (“X1” neoblasts). As a control, we inhibited *zfp-1* and found downregulation of *zfp-1* transcript as well as the epidermal (“zeta-neoblast”) marker *egr1* but not *soxP-1* or *hnf4,* as shown from prior studies (van Wolfswinkel et al., 2014) (Figure 3E). *mink1(RNAi)*, by contrast, did not significantly change the expression of any markers of specified states within X1 cells under conditions causing the impaired overall differentiation ability (Figure 3E). Therefore, *mink1* likely does not control specification within neoblasts, but rather the ability to produce post-mitotic differentiating cells.

### *Adenylate kinase-2 (ak2)* controls neoblast specification into intestine

Selective disruptions to individual lineages were rare in the screen, but one such target was *adenylate kinase 2* (*ak2*). *ak2* encodes a protein located in the mitochondrial membrane that catalyzes the reversible transfer of the terminal phosphate group between AMP and ATP in order to regulate energy homeostasis (Lagresle-Peyrou et al., 2009). *ak2* RNAi for 3 weeks (6 dsRNA feedings) led to failed regeneration (Figure 4A) and eventually lesions and lysis. qPCR expression data from the homeostasis screen revealed that *ak2(RNAi)* animals did not have impaired expression of neoblast markers (*smedwi-1, pcna, soxP-1, zfp-1*) or of *dd554+* pharynx progenitors or *nanos+* germline stem cells. However, the gamma-neoblast marker *nkx2.2* was significantly decreased, suggesting *ak2* could have a role in intestine formation (Figure 4B). To confirm this result, we used FISH to detect neoblasts (*smedwi-1*) and differentiate mature intestinal cells (*porcupine*) (Figure 4C). In *ak2(RNAi)* animals, the distribution of neoblasts was normal, but the intestine severely decayed. *ak2* activity could theoretically only be required for the maintenance and survival of mature intestinal cells, rather than their production. To test this possibility, we performed short term inhibition of *ak2* (4 dsRNA feedings, 17 days RNAi), then amputated animals to generate regenerating trunk fragments. Control animals produced new intestinal cells (marked by *porcupine*) by 8 days of regeneration, but *ak2(RNAi)* animals were unable to produce differentiated intestine cells (Figure 4D). Taken together, these results suggest *ak2* specifically promotes the production of the intestinal lineage in planarians rather than promoting neoblast self-renewal or only in maintaining mature intestine cells.

**Figure 4.**
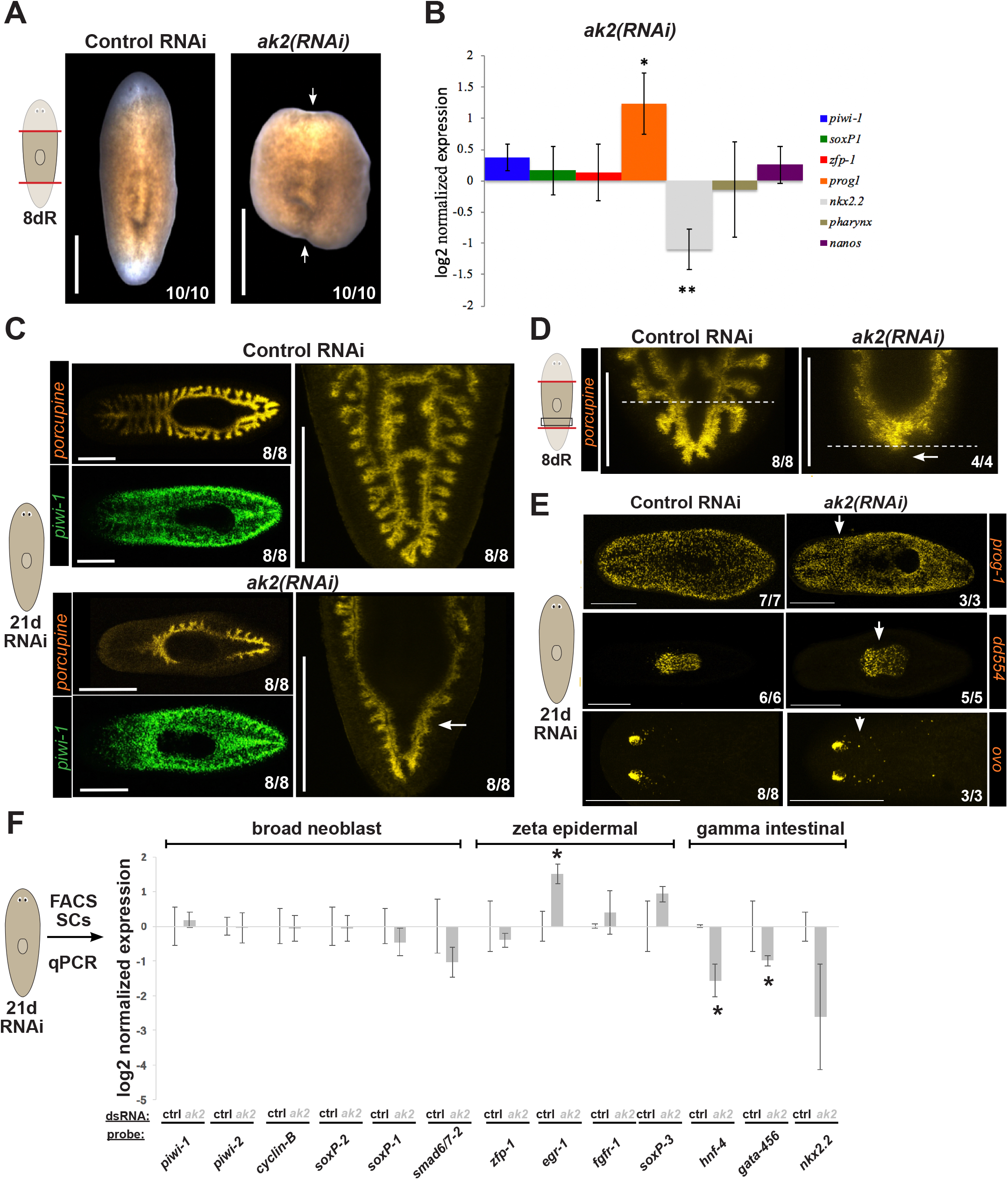
The energy metabolism kinase *ak2* is required for intestine specification. (A) Live animals scored at 8 days post-amputation showing that *ak2* RNAi prevented blastema formation (6 dsRNA feedings over 21 days). (B) Average log2 fold-change qPCR in whole-animal *ak2(RNAi)* normalized to control dsRNA conditions plotted from the primary screen in Figure 1C and Figure S2. Bars show averages of 3 biological replicate, error bars show standard deviations, asterisks indicate p<0.05 by unpaired two-tailed t-test versus control conditions. (C-E) In situ hybridizations of *ak2(RNAi)* versus control RNAi in uninjured animals (C, E) in regenerated trunk fragments 7 days after tail amputation (D), detecting neoblast marker *smedwi-1 (piwi-1)*, intestine marker *porcupine,* epidermal progenitor marker *prog-1*, eye progenitor marker *ovo* and pharynx progenitor marker *dd554* (6 dsRNA feedings over 21 days). Dotted lines indicate boundary between new/old tissue at the wound site in D. *ak2* RNAi caused reduction in intestine staining in homeostasis (C) and prevented formation of new intestine in regeneration (D) without reductions to *prog-1,* migratory *ovo+* eye progenitors, or pharynx progenitors (arrows) (E). (F) Planarian G2/S/M cells of the “X1” population were FACS sorted in biological replicate from control RNAi, and *ak2(RNAi)* animals, and markers of major neoblast subpopulations analyzed by qPCR. Bars show average log2 fold-change versus control RNAi conditions from 4 biological replicates obtained from 4 animals per replicate, error bars are standard deviations, and * denotes p < 0.05 by unpaired two-tailed t-tests versus control conditions. In *ak2(RNAi)* animals, markers of “gamma” neoblasts specifying toward intestine fate *hnf-4, gata4/5/6* and *nkx2.2* were downregulated in neoblasts without downregulation of pan-neoblast markers (“broad neoblast”), epidermal specifying neoblasts (“zeta”). Bars, 800 (A) or 300 microns (C-E).

We further examined the prediction from the qPCR and RNAseq that the role of *ak2* to promote differentiation was specific for intestine and not also other lineages. We tested this hypothesis by detecting the expression of progenitor cells of other lineages by FISH of the epidermal progenitor marker *prog-1*, the pharynx progenitor marker *dd554*, and the eye progenitor marker *ovo* (Figure 4E). In homeostatic animals, *ak2* inhibition did not lead to a visible increase or decrease of these progenitors. We next examined expression profiles of FACS-purified X1 neoblasts to determine whether *ak2* promotes the specification of intestinal precursor cells or is important for the maturation of this cell population after proliferative exit. In G2/S/M-purified neoblasts from *ak2(RNAi)* versus control RNAi animals profiled by qPCR, markers for intestine-specified neoblasts were downregulated (*hnf4*, *gata-4/5/6*) (Figure 4E). *nkx2.2* expression showed a stronger downregulation than the other two markers for this cell type, though we note that due to greater variability this was not statistically significant. By contrast, most other lineages remained unchanged after *ak2* RNAi, including pan-neoblasts (*gapdh, smedwi-1, smedwi-2, cyclinB),* sigma-neoblasts (*soxP-2, soxP-1, smad6/7*), zeta-neoblasts (*zfp-1, fgfr1, soxP3*) (Figure 4F). *ak2* RNAi caused a modest increase to *prog-1* expression levels in intact animals (Figure 4B) and also caused upregulation of the epidermal (“zeta”) neoblast marker *egr-1* within X1 cells, suggesting that in addition to promoting intestine regeneration, *ak2* could inhibit aspects of epidermal differentiation. Together these results are consistent with the hypothesis that *ak2* is required for specification of gamma-neoblasts, leading to the production of mature intestine cells. Due to the role of *ak2* in maintaining energy homeostasis, these results point could either suggest that intestine differentiation is secondarily regulated by global levels of ATP/ADP or perhaps the process of intestine differentiation is particularly sensitive to changes in ATP/ADP levels as regulated by *ak2*.

### *nat10* RNA acetylase facilitates epidermal progenitor maturation

Other tissue-specific regulators of differentiation were identified in the screen. *nat10* had RNAi caused an unusual profile of strongly elevated expression of epidermal progenitor markers and so is a candidate for control of epidermal differentiation either negatively or at a downstream step in progenitor maturation. *nat10* encodes an RNA N4-cytidine acetyltransferase that modifies rRNA and mRNA, and promotes translation (Arango et al., 2018). Following 6 feedings of *nat10* dsRNA, animals completely failed to regenerate (Figure 5A). A more detailed analysis of the expression phenotype revealed that *nat10(RNAi)* animals did not have changes to neoblast markers (*smedwi-1, soxP-1, zfp-1*) but instead had increases to epidermal lineage progenitors *prog-1* and *agat-1* (Figure 5B). Other differentiation markers of the pharynx (*dd554*) and germline stem cells (*nanos*) were not changed, and intestine differentiation marker *nkx2.2* was weakly and not statistically significantly downregulated. Therefore, *nat10* activity likely has a specific function in epidermal differentiation.

**Figure 5.**
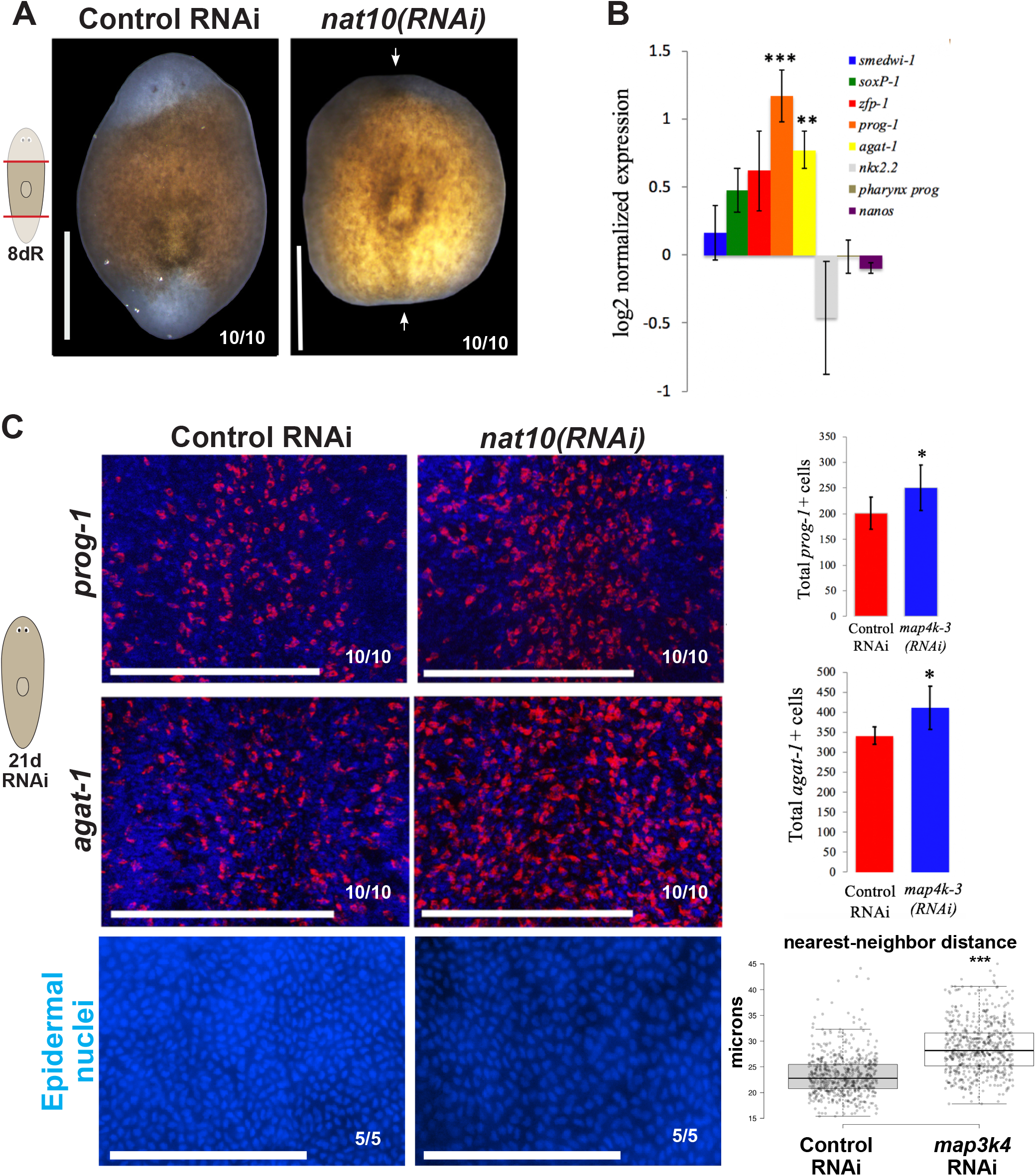
The RNA acetylase *nat10* limits abundance of epidermal progenitors. (A) Live phenotypes 8 days after amputation showing *nat10* RNAi prevented blastema formation after 6 dsRNA feedings over 21 days. (B) Bars show average log2 fold-change qPCR detection in whole-animal *nat10(RNAi)* versus control dsRNA, replotted from primary screen in Figure 1C and Figure S2. Error bars show standard deviations, and ** indicates p<0.01 from unpaired two-tailed t-tests. *nat10(RNAi)* caused upregulation of *prog-1* and *agat-1* markers of epidermal progenitors without significantly impacting other fate markers. (C) FISH of epidermal progenitor markers and Hoechst staining to visualize nuclei of mature epidermis layer after control or *nat10* homeostatic RNAi (6 dsRNA feedings over 21 days). Right graphs show number of cells per prepharyngeal field-of-view measured as either *prog-1* or *agat1+* (n=5 animals) or nearest-neighbor distances between nuclei (n=5 animals). * indicates p<0.05 by unpaired two-tailed t-tests. Nearest neighbor distance computed for epidermal nuclei in fields of view taken in a prepharyngeal region for each of 5 animals per condition, *** indicates p<0.001 by unpaired two-tailed t-tests. Bars, 800 (A) or 300 microns (C).

In order to investigate whether *nat10* inhibits the abundance of epidermal cell progenitors, animals were fed with control or *nat10* dsRNA 6 times and FISH was performed to detect either *prog-1* or *agat-1* mRNA (Figure 5C). Quantification of the number of *prog-1* or *agat-1* positive cells showed a significant increase in the abundance of these cells in *nat10(RNAi)* versus control RNAi conditions. Therefore, *nat10* inhibition leads to greater numbers of *prog-1+* and *agat-1+* cells rather than only changing their expression levels per cell. We considered two possible explanations for these observations, that either *nat10* negatively regulates epidermis formation or is critical for epidermal progenitor maturation to terminal differentiation. Because *nat10(RNAi)* animals fail to regenerate altogether, we deemed this second possibility more likely. Consistent with this hypothesis, nuclei within the mature epidermis of *nat10(RNAi)* animals had a decreased density, a condition known to arise from impaired differentiation of new epidermal cells (van Wolfswinkel et al., 2014). Taken together, these results suggest *nat10* limits the abundance of epidermal progenitors without altering neoblasts or mature epidermis, we suggest through ensuring their normal maturation.

### *pak-1* kinase restricts neoblast localization and prevents outgrowth of dorsal tissue

Phenotypes of neoblast over-proliferation have only rarely been described, so we were interested in uncovering genes limiting expression of neoblast markers (Gonzalez-Estevez et al., 2012; Oviedo et al., 2008). Phenotypes of excess neoblast marker expression were also rare in the screen, and had low effect sizes. Of all factors screened, *pak1(RNAi)* uniquely caused a reproducible but relatively small upregulation of both *piwi-1* and *pcna,* so we further examined these animals for additional phenotypes. *pak1* encodes a conserved kinase and critical effector linking Rho family GTPases to cytoskeleton reorganization and nuclear signaling (Kumar et al., 2009; Kumar and Vadlamudi, 2002; Vadlamudi and Kumar, 2003). Pak1 factors are targets for small GTP binding proteins Cdc42 and Rac and have roles in a wide range of biological activities such as cell migration, motility, and morphology (Brown et al., 1996; Sells et al., 1997; Zhao et al., 1998). In planarians, *pak1* inhibition strongly reduced blastema formation and resulted in midline indentations (Figure 6A). After homeostatic inhibition, *pak1(RNAi)* animals had small but significant increases to all neoblast markers we tested, as well as early progenitors *prog-1* and *agat-1* (Figure 6B). However, at this timepoint (6 dsRNA feedings, 21 days), there were no obvious signs of neoblast over-proliferation such as tissue outgrowths.

**Figure 6.**
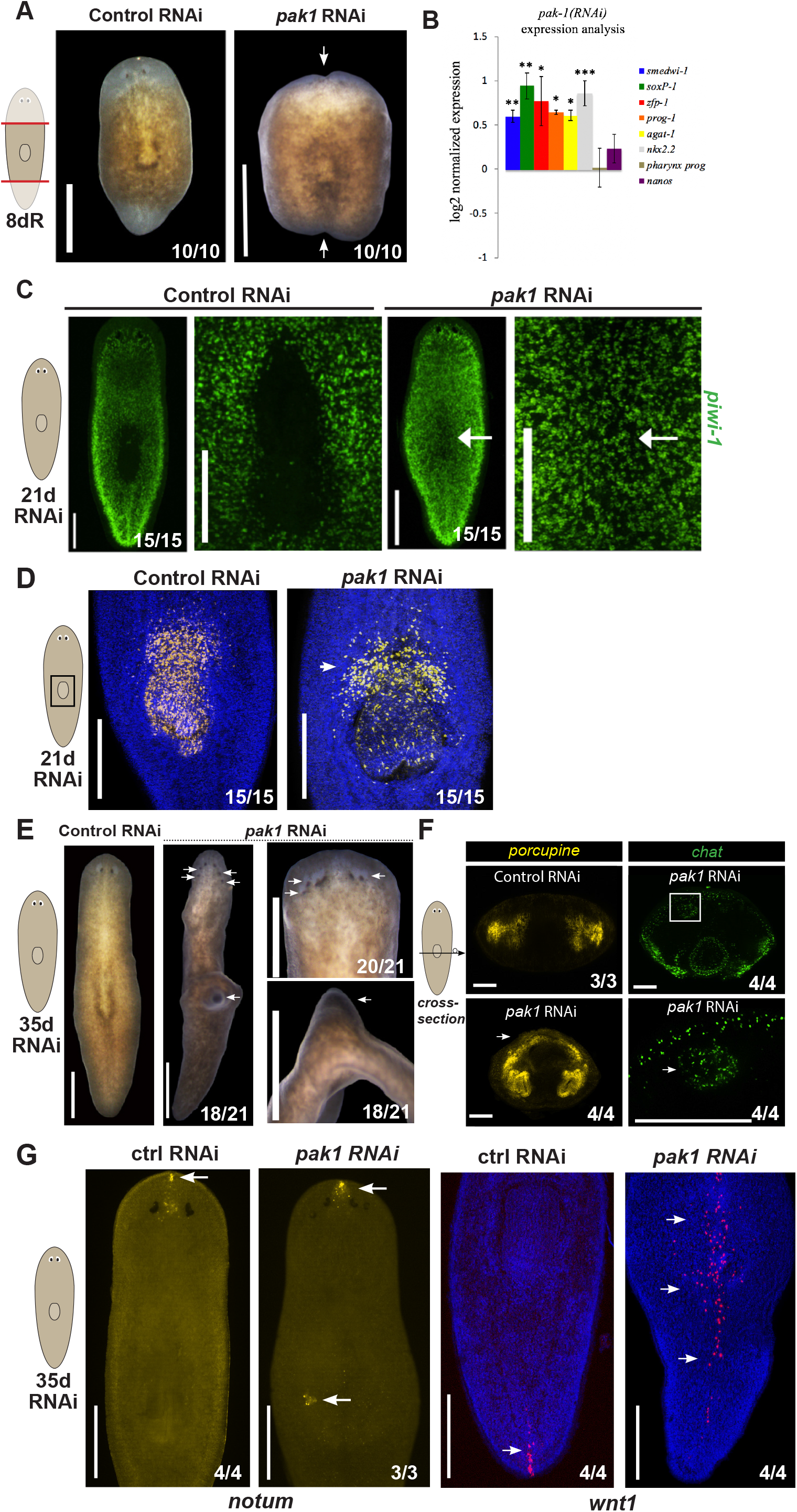
*pak1* inhibition mispatterns the body, mistargets neoblasts, and causes outgrowths. (A) Live phenotypes showing *pak1(RNAi)* animals regenerate with small, indented blastemas. (B) Average log2 fold-change qPCR in whole-animal *pak1(RNAi)* versus control dsRNA from primary screen (Figure 1C, Figure S2). Error bars show standard deviations. *pak1* RNAi caused upregulation of several neoblast and progenitor markers (*smedwi-1/piwi-1, soxP-1, zfp-1, prog1, agat-1*). (C-D) Homeostatic *pak1(RNAi)* and control RNAi animals stained by FISH to detect *piwi-1* and *dd554* pharynx progenitor marker after 6 dsRNA feedings over 21 days. (C) *pak1* RNAi caused *piwi-1* neoblasts to localize to the pharynx region normally devoid of neoblasts (arrows), (D) and caused *dd554+* pharynx progenitors to mislocalize anteriorly (arrow). Right panels in each treatment show enlargements of pharynx region. (E) Long-term inhibition after 12 dsRNA feedings over 35 days *pak1* caused formation of ectopic eyes posteriorly in the head, and a dorsal outgrowth at the position of the pharynx (arrows). Right, insets show enlargement of the anterior (top panel, dorsal view) or trunk region of animals (bottom panel, side view). (F) Cross-sections of animals stained by FISH to detect gut branches (*porcupine*) and neurons (*chat*). *pak1* RNAi caused ectopic gut branches at the location of the dorsal trunk outgrowth (arrows), as well as ectopic *chat+* neural clusters dorsally (bottom right panel shows enlarged inset from region of box in top right panel). (G) FISH of *notum* and *wnt1* in homeostatic *pak1* RNAi versus control RNAi. *pak1(RNAi)* animals had an intact *notum+* anterior pole and dorsal trunk outgrowths frequently had a cluster of *notum+* cells suggestive of this tissue displaying anterior identity (arrows). *pak1(RNAi)* caused anterior and midline expansion of the population of *wnt1+* posterior pole cells (arrows). Bars, 800 (A), 300 (C), or 150 microns (F).

We stained animals homeostatically inhibited for *pak1* after 21 days of RNAi (6 feedings) and found that *piwi-1+* cells invaded a region near the pharynx normally devoid of neoblasts (Figure 6C). In concert, the pharynx progenitor marker *dd554* became dispersed and localized further anterior, consistent with a progressive modification to the trunk regionalization as displayed by altered neoblast distributions (Figure 6D). *pak1*(RNAi) homeostatic animals ultimately displayed a range of patterning defects after 35 days of inhibition (12 dsRNA feedings), including extra sets of eyes located posterior to the original eyes, and a dorsal outgrowth at the location of the region invaded by surrounding neoblasts (Figure 6E). FISH stained animals sectioned through the trunk region of *pak1(RNAi)* animals revealed that the outgrowth contained ectopic *porcupine+* intestine tissue as well as ectopic *chat+* neurospheres aggregates (Figure 6F) similar to those found from inhibition of *integrin-beta-1*, in which neoblast migratory targeting is impaired (Bonar and Petersen, 2017; Seebeck et al., 2017). Given the overt patterning defects after *pak1* inhibition, we probed whether these body transformations were coincident with alterations to signals controlling planarian body-wide patterning. The outgrowths frequently included a focus of *notum+* cells similar to a *notum+* region at the anterior pole organizing center of normal animals (Figure 6G) (Petersen and Reddien, 2011; Vasquez-Doorman and Petersen, 2014). However, expression of *notum* at the anterior pole of was unaffected by *pak1* inhibition. By contrast, the master regulator of posterior formation, *wnt1* (Petersen and Reddien, 2009; Schad and Petersen, 2020), was dramatically expanded along the midline of *pak1(RNAi)* animals. We suggest that *pak1* acts as a patterning factor, such that *pak1* RNAi modifies the body plan and leads to a body transformation enabling aberrant targeting of neoblasts to these territories, followed by inappropriate outgrowth in this region. Taken together, these results identify a new role control point regulating neoblast distribution across the body important for preventing tissue outgrowths.

## DISCUSSION

We present a new strategy for uncovering phenotypes in planarians and report new functions for a variety of factors in regulating tissue homeostasis. We performed a high-throughput screen of hundreds of kinases and other signaling molecules and found 66 genes required for homeostasis and regeneration. In order to uncover phenotypes of neoblast dysfunction, we developed a high-throughput RNA extraction method and performed qPCR on markers of different neoblast classes and progenitors of multiple cell lineages. This tool allowed us to distinguish between a variety of neoblast regulators including promoters of self-renewal, differentiation, the intestine lineage, as well as inhibitors of neoblasts and their progenitors. The spectrum of changes to key cell state markers in whole animals directly permitted factors to be assigned to particular functions in the planarian stem cell homeostasis hierarchy (Figure 7).

**Figure 7.**
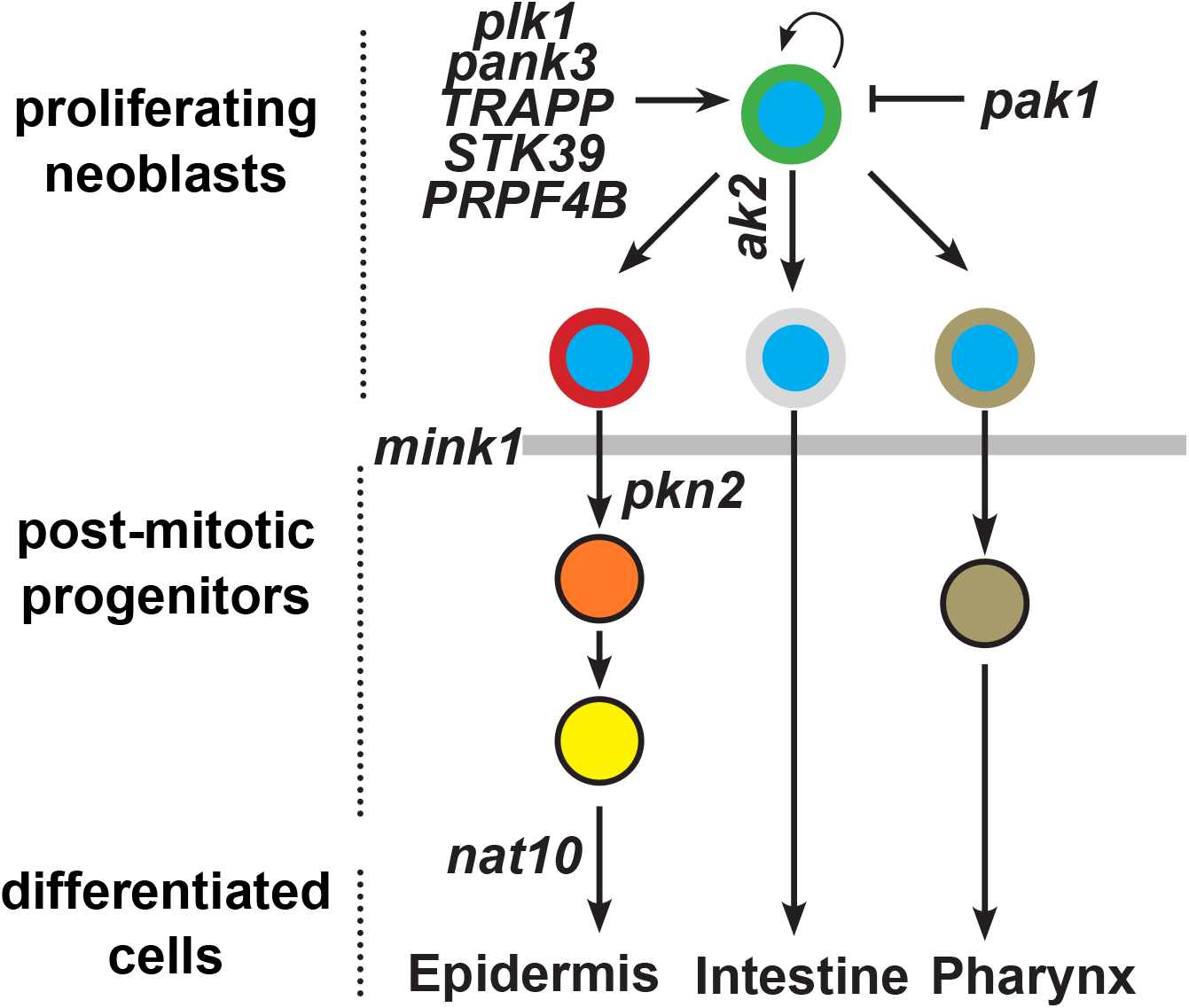
Model placing discovered genes into pathway for tissue homeostasis by neoblasts. Neoblasts (blue circles) self-renew and generate all differentiated cells of adult planarians. Specification occurs within the neoblast population (outline colors), followed by terminal differentiation, either directly after cell cycle exit or through post-mitotic progenitor intermediates. Several genes were identified as critical for maintenance of neoblasts and are likely important for self-renewal of these cells (eg., *plk1*, *pank3*, *trrap*, *stk39*, etc). *mink* kinase is critical for exit from the proliferative neoblast state (shaded box). *ak2* controls intestine differentiation at the level of early neoblast specification. *nat10* limits progenitor numbers likely through control of their maturation to terminal differentiation. *pak1* limits neoblast distribution in the body and controls patterning to ensure appropriate homeostasis.

Although the approach was extensively validated by histology with several targets, there could be limitations to the strategy of expression profiling during RNAi screening. In order to maximize the number of targets analyzed, we restricted our analysis to a single timepoint during the progression of each phenotype, which could in principle to false negatives or incomplete information of cell states. For example, in the qPCR secondary screen, RNAi of *smg-1*, a factor whose inhibition can result in elevated neoblast abundance (Gonzalez-Estevez et al., 2012), was surprisingly detected as a positive regulator of neoblast gene expression (Figure 1C). Some planarian neoblast reduction phenotypes can pass through an initial state of elevated neoblasts, perhaps because of growth arrest prior to depletion, for example in *p53* RNAi or *piwi-2* RNAi (Pearson and Sanchez Alvarado, 2010) (Reddien et al., 2005b). Our results suggest that neoblast depletion is a possible outcome from long-term *smg-1* inhibition, and we speculate this loss of neoblast expression could occur via a similar mechanism. In addition, our analysis revealed an expected correspondence between expression changes of the early versus late epidermal progenitor markers *prog-1* and *agat-1* were observed after knockdown of several control and experimental conditions (*repA1*, *plk1*, *ANAPC1*, *bruli*, *zfp-1*, *akt*, *mink1*). However, some treatments caused downregulation of *prog-1* but not *AGAT-1* (*papss2, mnat1, pank3, ank1, cdk12-2, lamtor, csnk2a-1, trrap, ppip5k2, pkn2*). Given that lethal irradiation causes *prog-1* downregulation prior to downregulation of *AGAT-1* expression (Eisenhoffer et al., 2008), a likely explanation is that some phenotype states detected in the screen represented epidermal differentiation dysfunction that had not yet progressed to affect later-stage progenitors.

Our analysis revealed several categories of phenotypes involved in planarian stem cell activities. The largest phenotypic category identified were factors required for neoblast abundance as well as production of differentiating progenitors. Genes in this category carry out a variety of functions including the regulation of cell proliferation (*Rb, plk1, cdk12, ANAPC1, RepA1),* transcriptional regulation (*mnat, cdk7*), cell-signaling, and other processes (*ank1*, *pank3*). By contrast, *mink1* represented a class of factor required strongly for producing post-mitotic differentiated cells across lineages. The rarity of such factors suggests that the process of neoblast proliferative exit is a relatively specific control point likely of key importance for understanding the biology of planarian pluripotent cells.

Factors promoting specific lineages were also rare, with a small number of newly discovered factors specifically important for epidermis (*pkn2*, *akt*) and intestine (*ak2*), and none specifically required for other lineages. No factors were identified as specifically required for *dd554*+ pharynx progenitor cells, and RNAseq of many conditions also revealed additional categories of losses of specific progenitor types. The lack of phenotypes selective for failed differentiation across other cell lineages could perhaps arise from the strategy to perform expression profiling only of factors required for overall regeneration and homeostasis in the primary screen. In addition, most other lineages in the animal (i.e., eyes, protonephridia, most neural cell types, etc) are thought to lack a progenitor state that can be uniquely detected in whole animals by expression of single genes. The importance of an energy metabolism factor *ak2* specifically for intestinal differentiation suggests the possibility that global energy levels feed back onto production of tissues involved in absorption and/or energy storage. The intestine has previously been suggested to act as a neoblast niche due to the fact that neoblast reside nearby intestine cells throughout planarians (Forsthoefel et al., 2012), and the intestine produces apolipoproteins that control the process of neoblast differentiation (Wong et al., 2022).

Finally, the methodology allowed identification of novel classes of dysfunction in neoblast-dependent tissue homeostasis. *nat10* inhibition affected a step in epidermal differentiation leading to a novel defect in which epidermal progenitor cells accumulated. *pak1* RNAi led to increased expression of neoblast markers and a novel phenotype of neoblast mislocalization within the pharynx area and subsequent mispattered outgrowths. These results indicate that expression profiling as a strategy for phenotyping in screens can identify unexpected control points.

The process of tissue homeostasis and its role in whole-body regeneration is likely to be complex given the large set of cell types necessary to generate and their appropriate targeting. Planarian neoblasts represent a powerful model for studying the mechanisms of self-renewal, specification, and differentiation in adult stem cells during homeostasis and regeneration. Our study offers a new method of uncovering genes specifically regulating these processes in planarians, allowing for an unbiased screening to discover new regulators and new control points in complex developmental regulatory hierarchies.

## Acknowledgements

We thank members of the Petersen lab for critical comments and support. C.P.P. acknowledges funding from the National Institutes of Health, USA (NIGMS R01GM129339 and R01GM130835), and pilot project funding from the NSF-Simons Center for Quantitative Biology at Northwestern University, an NSF (1764421)-Simons/SFARI (597491-RWC) MathBioSys Research Center.

## Author Contributions

E.G.S and C.P. designed the study. E.G.S performed the experiments, and E.G.S. and C.P. wrote the paper.

## MATERIALS AND METHODS

### Planarian culture

Asexual *Schmidtea mediterranea* animals (CIW4 strain) were maintained in 1x Montjuic salts between 18–20°C. Animals were fed a puree of beef liver and starved for at least 5 days before experiments.

### Fluorescent in situ hybridization (FISH)

Animals were fixed and bleached as described previously (King and Newmark, 2013; Pearson et al., 2009). Riboprobes (digoxigenin- or fluorescein-labeled) were synthesized by in vitro transcription (King and Newmark, 2013; Pearson et al., 2009). Antibodies were used in MABT/10% horse serum/10% Western Blocking Reagent (Sigma-Roche) at a concentration of 1:2000 for anti-digoxigenin-POD (Sigma-Roche #11207733910 RRID: AB_514500), and 1:2000 for anti-fluorescein-POD (Sigma-Roche #11426346910 RRID: AB_840257). For multiplex FISH, peroxidase conjugated enzyme activity was quenched between tyramide reactions by sodium azide treatment (100 mM in 1xPBSTx) for 45 min at room temperature. Nuclear counterstaining was performed using Hoechst 33342 (Invitrogen, 1:1000 in 1x PBSTx).

### RNAi administration

Genes were either cloned by RT-PCR using primer sequences indicated in Table S1 or generated from cDNA clones from previously described *Schmidtea mediterranea* cDNA libraries as indicated in Table S1 (Sanchez Alvarado et al., 2002). RNAi by feeding was performed in vitro transcribed dsRNA mixed with 80% liver paste and 5% red food dye (Rouhana et al., 2013). In all RNAi experiments animals were fed RNAi food every 2-3 days for the length of experiment indicated. Animals were kept on 1% gentamicin in planaria water during dsRNA feedings. Plastic dishes were replaced each day before feeding and the day following feeding.

### High throughput RNA extraction

Animals were sorted into racked 96-well tubes (ClavePak 1158R63, Denville), using 3-4 biological replicates of 3-4 intact or regenerating fragments per replicate in ∼100ul of planarian media (1x Montjuic salts). Planaria media was replaced with 200 ul TEN buffer (0.1 M Tris-Cl pH 8.0, 0.01 M EDTA pH 8.0, 1 M NaCl) supplemented with SDS (0.4%) proteinase K (0.48 mg/ml) and beta-mercaptoethanol (0.14%) and pipetted gently with a multichannel pipette every 2-3 minutes at room temperature until homogenized (∼10 minutes), followed by addition of 600ul Trizol-LS with mixing, followed by 200 ul chloroform, mixing vigorously by pipetting then incubated at room temperature for 2-3 minutes. Plates of 96-well racked tubes were centrifuged at 4800 rpm at 4°C for 15 minutes on a Thermo ST16R table-top centrifuge with acceleration/deceleration setting 9. 300-350ul of aqueous layer was removed using multichannel pipettes, mixed with 600ul 70% ethanol and purified using 96-well column silica-membrane filter plates (Epoch Life sciences 2020-001) positioned over 2ml deep-well 96-well plates (VWR 40002-014) to collect flow-through, and centrifuged at 2300g, 4°C, for 5 minutes. Columns were washed with 700µL 3M Na Acetate, twice with 500µL 70% EtOH, and dried by using the same centrifugation parameters. Samples were eluted in 30ul MiliQ-purified H2O, spot-verified by nanodrop and Agilent Bioanalyzer pico assay and stored at -80C.

### High-throughput qPCR

cDNAs were synthesized using MultiScribe MMLV reverse transcriptase (Life Technologies) and oligo-dT (Qiagen) for 1 hour at 37°C. Primer sets for qPCR are described in Table S5. High-throughput qPCR reactions were assembled in 384-well plates using an TTP Labtech Mosquito Crystal and EvaGreen qPCR mastermix (Midwest Scientific BEQPCR-S) and reactions performed on a BioRad CFX384 in the Northwestern University High Throughput Analysis laboratory. Per-sample log2-fold change expression were determined by the delta-delta Ct method, normalizing to expression of *clathrin* probe set control and to within-batch RNAi negative controls targeting *Photinus pyralis* luciferase. Statistical significance was determined through unpaired two-tailed t-tests comparing ddCt values between control RNAi and target RNAi conditions. p-values were further corrected for false-discovery through the Benjamini-Hochberg method across conditions within the set of measurements obtained for each probe set. Log2 fold-change, standard deviations, p-values and false discovery adjusted p-values described in Table S2.

### RNA-seq

96-well RNAseq libraries were prepared using the Kapa stranded mRNA seq system and sequenced on a NextSeq500 using 75bp single-reads at the Duke University Center for Genomics and Computational Biology. Reads were aligned to the planarian transcriptome using bowtie1.2 and counts determined using HTseq, followed by testing for differential gene expression using DEseq in R. Full dataset with log2 fold-change and false discovery adjusted p-values is shown in Table S4 and data used in Figure 2 and Figure S6 is shown in Table S3. Sequences were deposited at NCBI as GEO entry GSE212136.

### Imaging/Quantification of images

Live images of animals were performed with a Leica MZ10F dissecting scope with a Leica DFC295 camera. FISH images were performed with a Leica DM5500B compound microscope with optical sectioning by Optigrid structured illumination, a Leica TCS SPE, or a Leica Stellaris 5 confocal compound microscope with mosaic scanning and stitching. Fluorescent images collected by compound microscopy are maximum projections of a z-stack and adjusted for brightness and contrast using Metamorph, Leica LAS, or Adobe Photoshop. Nuclei counting in Figure 5C was conducted in ImageJ using maximum projections of the 15-micron thick most outer Hoechst-stained tissue from field-of-view images of the dorsal prepharyngeal region to segment and quantify nuclei followed by calculation of nearest neighbor distance in 2-dimensions tabulated across 5 biological replicate samples.

### Cell Counting

In Figure 5C, *prog1+* and *agat1+* cells were imaged using a Leica DM5500B compound microscope with optical sectioning equipped with an Optigrid structured illumination device. After a maximum projection of z-stacks, cells were counted manually in Metamorph. Cell numbers were averaged and significance was determined by unpaired two-tailed t-tests.

## Supplemental figure legends

**Figure S1.**
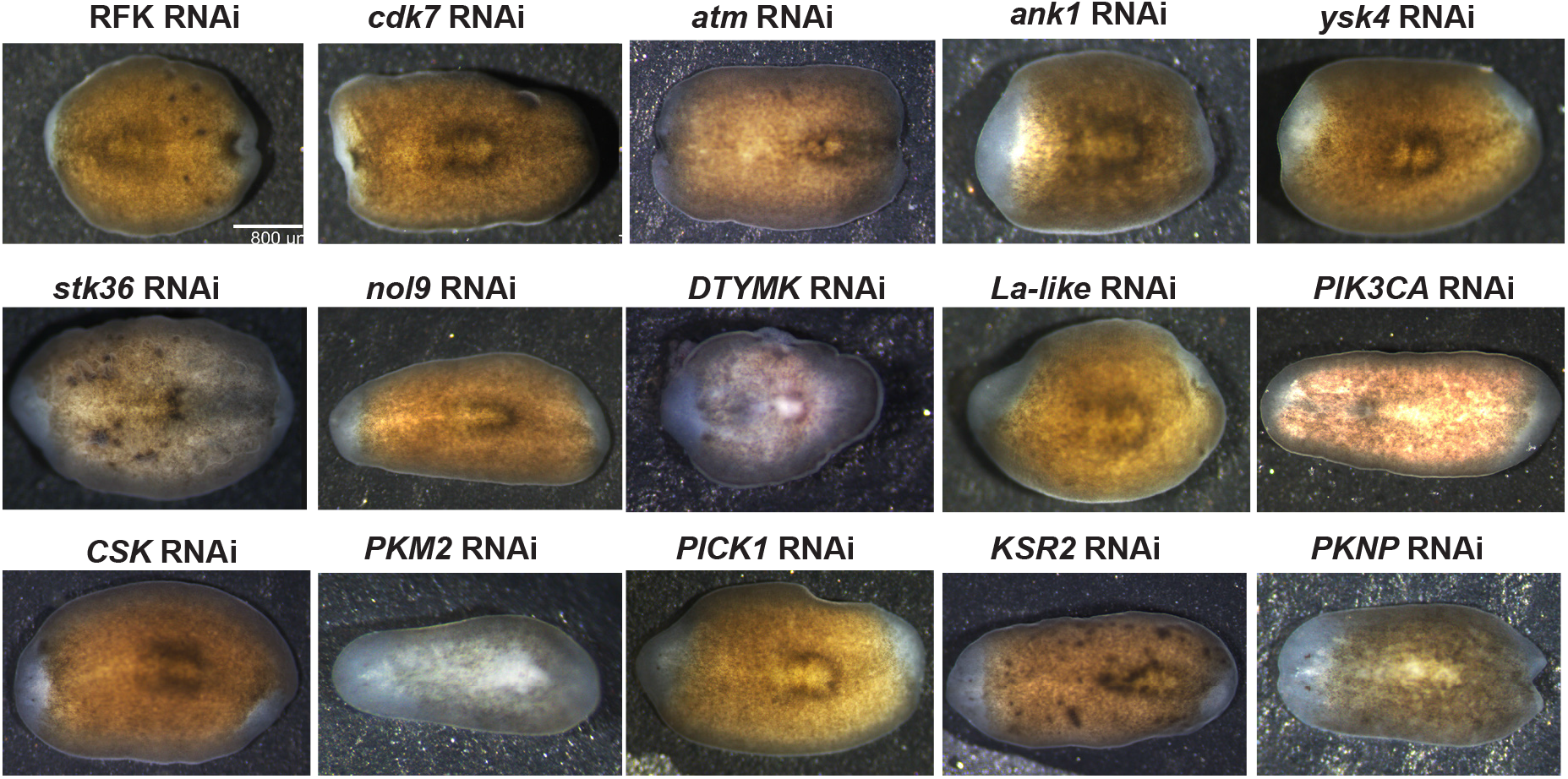
Live images of regeneration-deficient phenotypes recovered from the primary screen and imaged at day 8 during regeneration of a new head and tail from amputated trunk fragments to visualize blastema outgrowth. Additional details and comments in Table S1. Images represent 12/12 animals probed unless indicated otherwise in Table S1.

**Figure S2.**
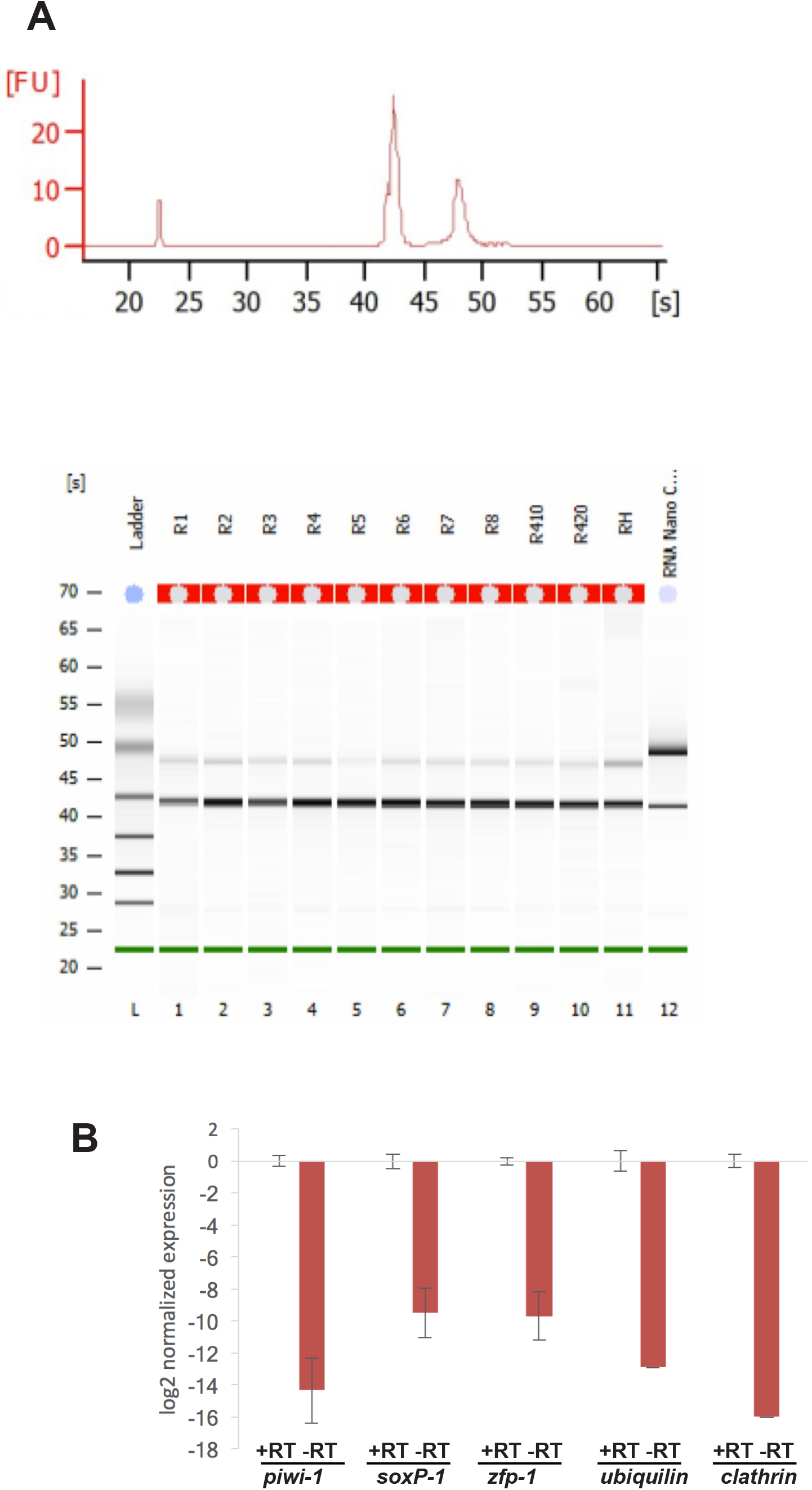
(A) PicoAnalyzer chip results from SDS/beta-mercaptoethanol/proteinaseK + Trizol RNA extraction method showing (top) representative absorbance trace and (bottom) virtual gel from multiple lanes (lanes 1-11 indicate total RNA from separate biological replicates of 4 animals per replicate, L is the ladder and lane 12 is positive control RNA from HeLa cells). (B) qPCR controls showing detection of mRNA and not genomic DNA. log2 expression for each probe-set comparing conditions of inclusion (+RT) or exclusion (-RT) of reverse transcriptase enzyme prior to qPCR. Bars show log2 average expression normalized to +RT condition from 3 biological replicates after flooring any samples lacking detection to Ct=40, and error bars show standard deviations. qPCR signal in the absence of reverse transcriptase was at least 10-fold lower for each probe.

**Figure S3.**
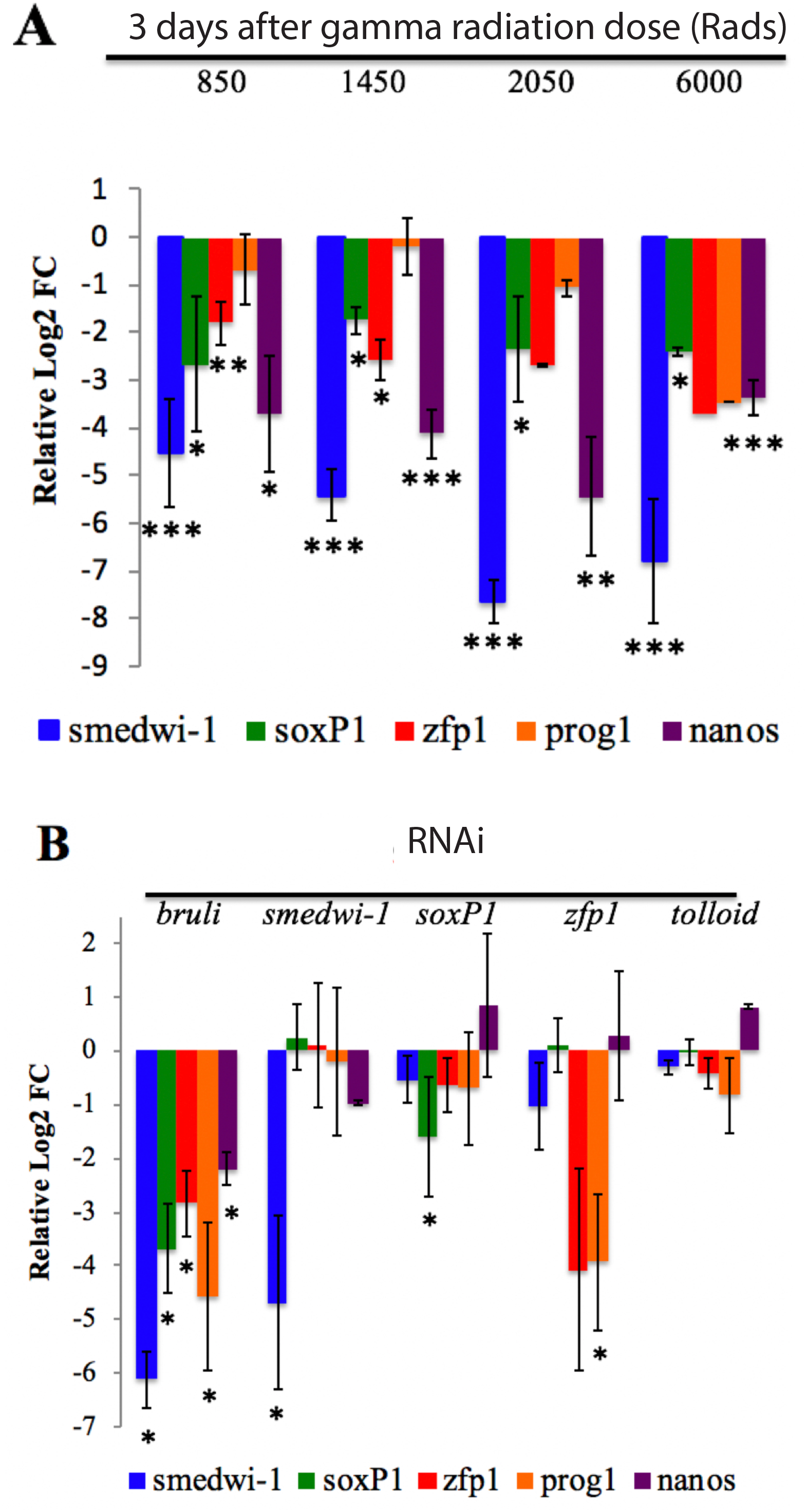
(A) Samples were subjected to a series of ionizing radiation and 3 days of homeostasis prior to RNA extraction and testing for expression of key neoblast/progenitor markers (neoblast markers *smedwi-1/piwi-1* and *soxP-1,* epidermal-specified neoblast marker *zfp-1*, epidermal progenitor marker *prog-1,* germline progenitor marker *nanos*). Graph shows average log2 fold-change expression values from 3 biological replicates normalized by the delta-delta Ct method to unirradiated animals and the housekeeping mRNA control *clathrin.* (B) qPCR as in (A) using samples inhibited by RNAi for the indicated genes then probed for expression of neoblast/progenitor markers. Graph shows average log2 fold-change expression values from 3 biological replicates normalized by the delta-delta Ct method to animals treated with control non-targeting dsRNA from *Photinus pyralis luciferase* and the housekeeping mRNA control *clathrin.* (A-B) Error bars show standard deviations, *p<0.05, **p<0.01, ***p<0.001, unpaired two-tailed t-tests.

**Figure S4.**
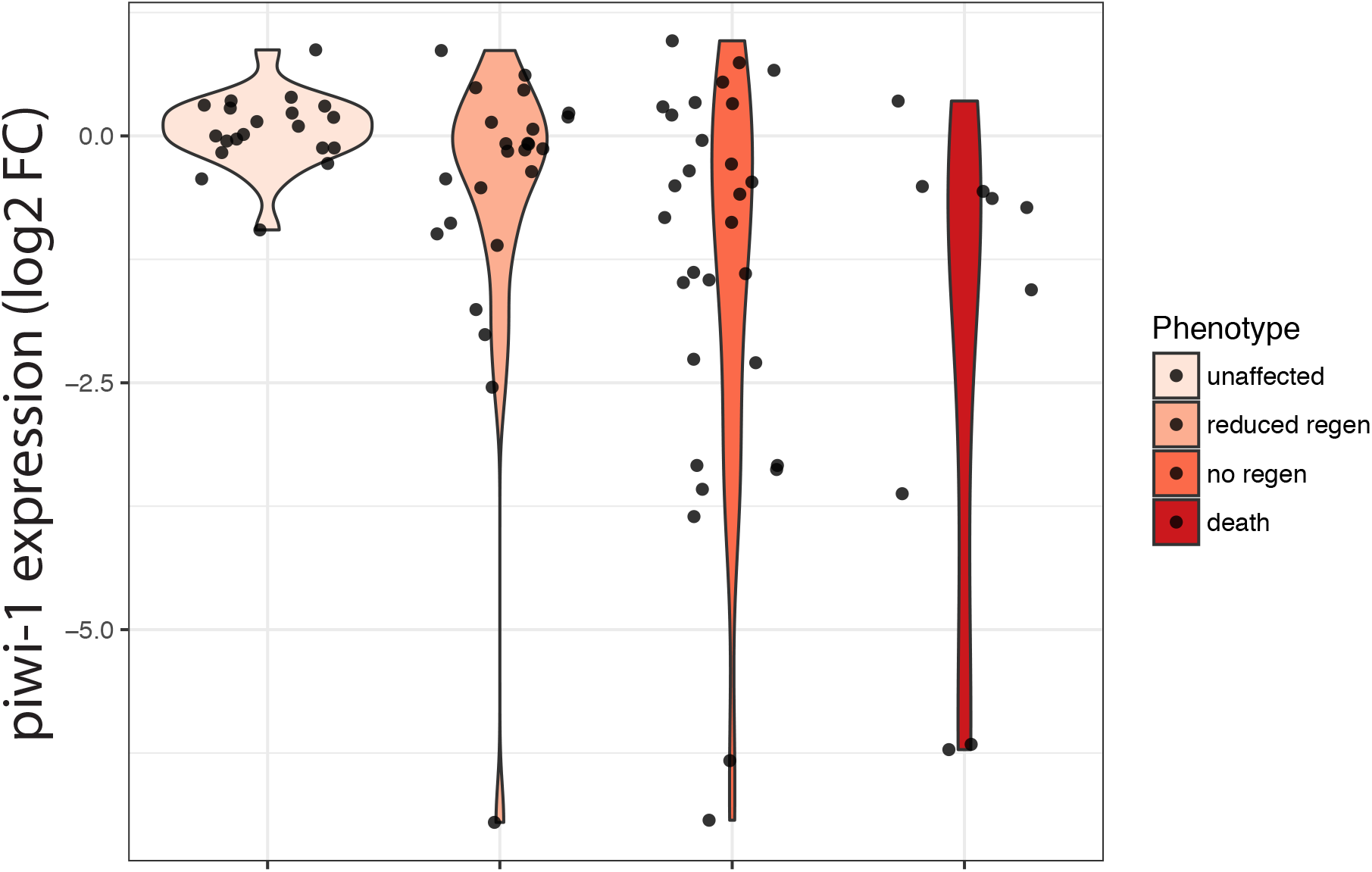
Relationship between phenotype strength as scored in the primary screen versus change in *piwi-1* measured in the secondary screen. Phenotypes are arranged in increasing severity (unaffected indicates normal regeneration, reduced regeneration indicates formation of reduced sized blastema, no regeneration indicates absence of blastema, death indicates animal lysis). In each category of regeneration or viability reduction, some treatments caused decreased expression of *piwi-1* while others did not. Therefore, factors responsible for self-renewal constitute a specific subset of those important for regeneration and viability.

**Figure S5.**
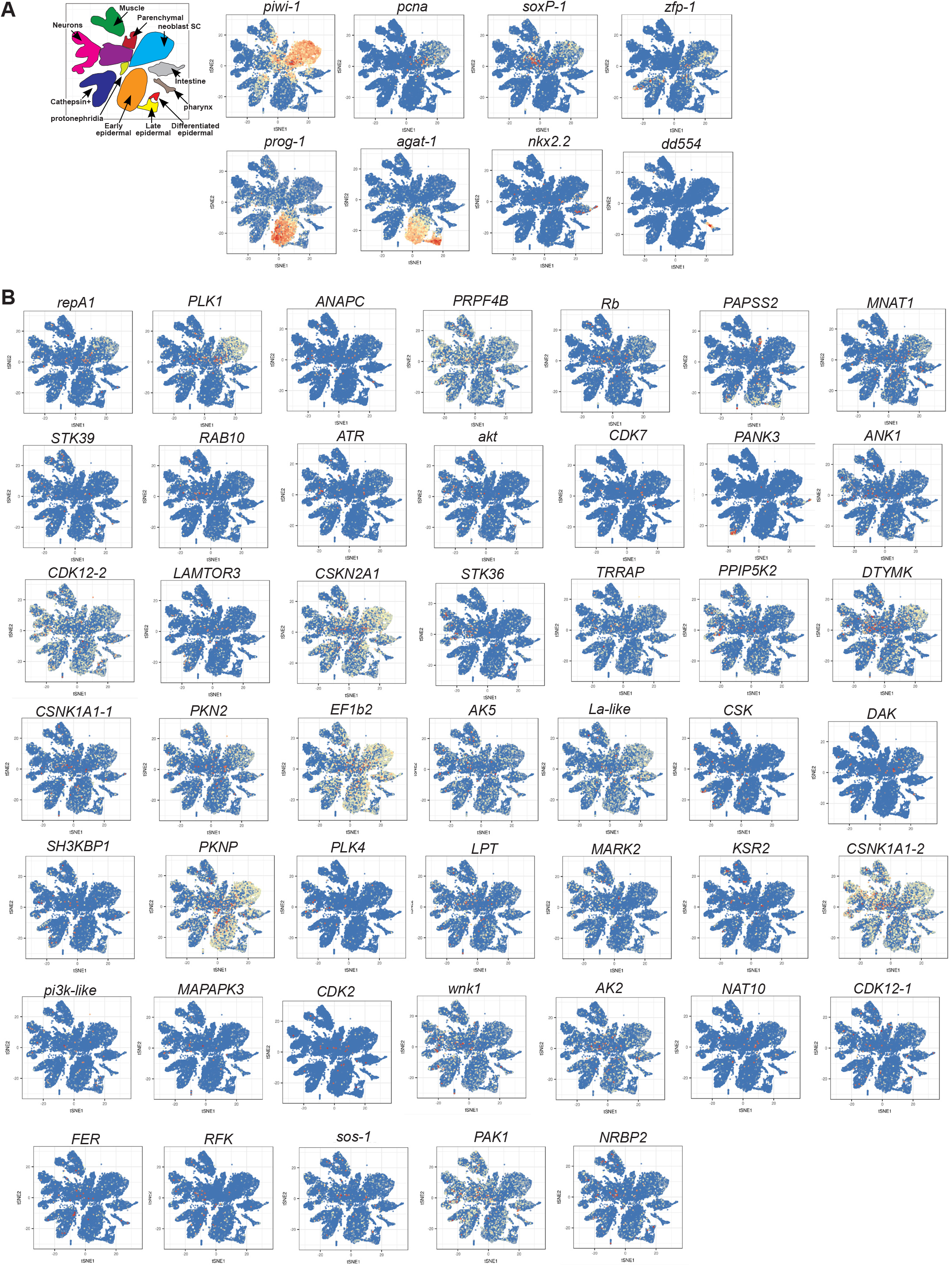
scRNAseq expression patterns of (A) markers used in qPCR screen and (B) genes recovered from the screen, obtained from planarian cell atlas (Fincher et al., 2018) (digiworm.wi.mit.edu). Legend indicates identities of major clusters. Genes in (B) are listed in the order they are presented in the heatmap in Figure 1C. Most factors analyzed have broad expression across multiple cell types, including in neoblasts. Therefore, a majority of factors recovered from the screen as important for neoblast maintenance or early differentiation could either operate autonomously in neoblasts or nonautonomously for their function. For example, factors important for self-renewal (*mnat1, cdk7, papss2, prpf4b, cdk12, ank1*) have expression in neoblasts and also in most cases in multiple types of differentiated cells, and these factors were required for neoblast maintenance. Some recovered factors are expressed specifically in neoblasts (*plk1*, *cdk2*). *pkn2* has strong expression in neoblasts as well as several differentiated cell types. *ak2*, *nat10*, and *pak1* have broad expression. *pank3* is expressed primarily in subsets of intestine and Cathepsin+ cells, yet is required for neoblast maintenance, suggesting this factor could facilitate the neoblast microenvironment as a potential niche factor.

**Figure S6.**
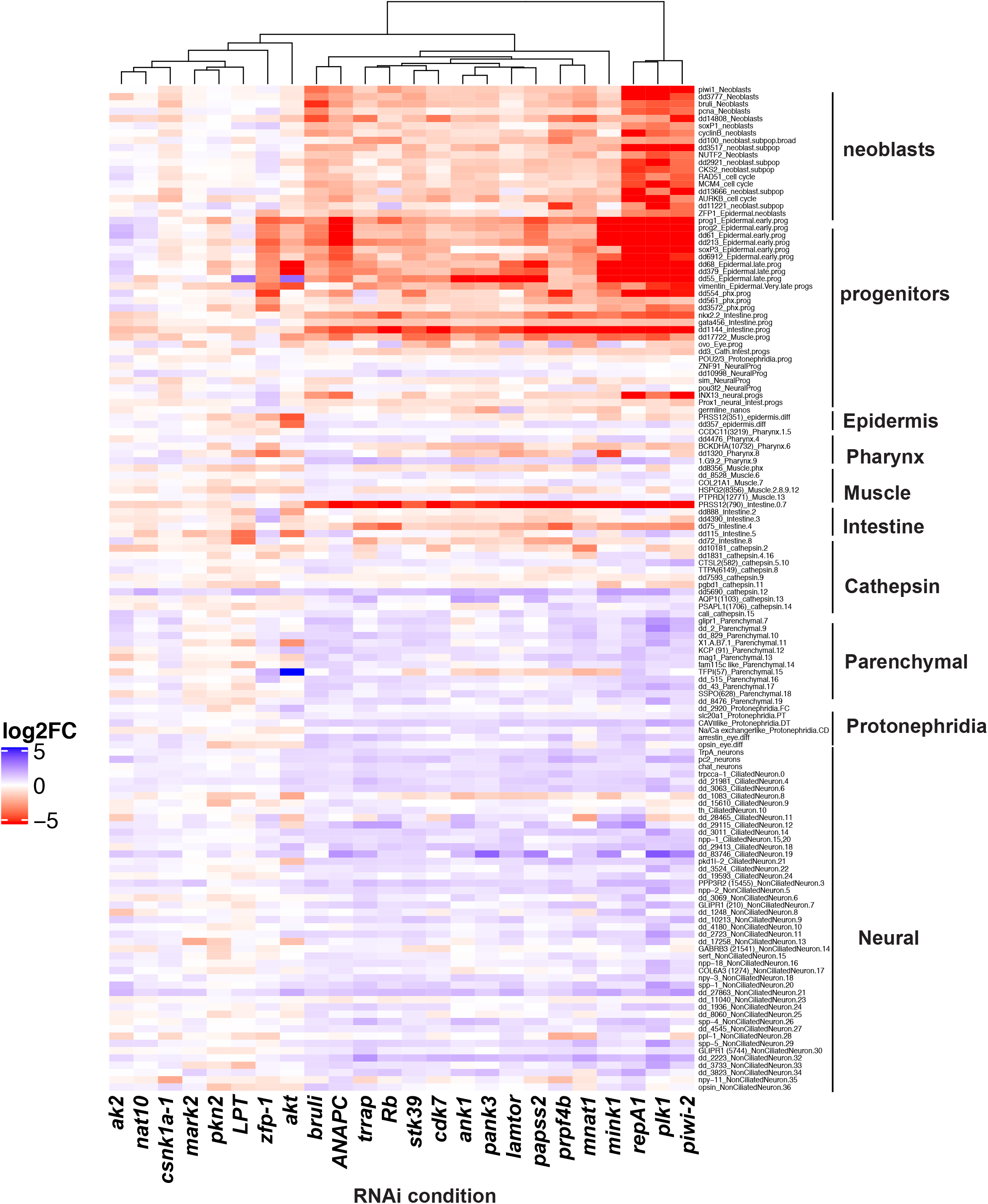
RNAseq was performed on 24 of the RNAi conditions used in the secondary screen. Average log2 fold-change expression values are shown for several known neoblast markers and markers of major progenitor and differentiated cell types in the animal described in the scRNAseq planarian cell atlas. Heatmap shows log2 fold-change of 3 biological replicates compared to negative Control RNAi conditions targeting the *Photinus pyralis* luciferase gene not present in the planarian genome. Columns show RNAi treatments, rows show marker gene expression following RNAi. Full dataset expression log2 fold-change expression values, and adjusted p-values shown in Table S3. A majority of tested conditions resulted in reduced expression of most neoblast and/or progenitor markers, especially for fast-turnover tissues such as epidermis, intestine, and pharynx.

**Figure S7.**
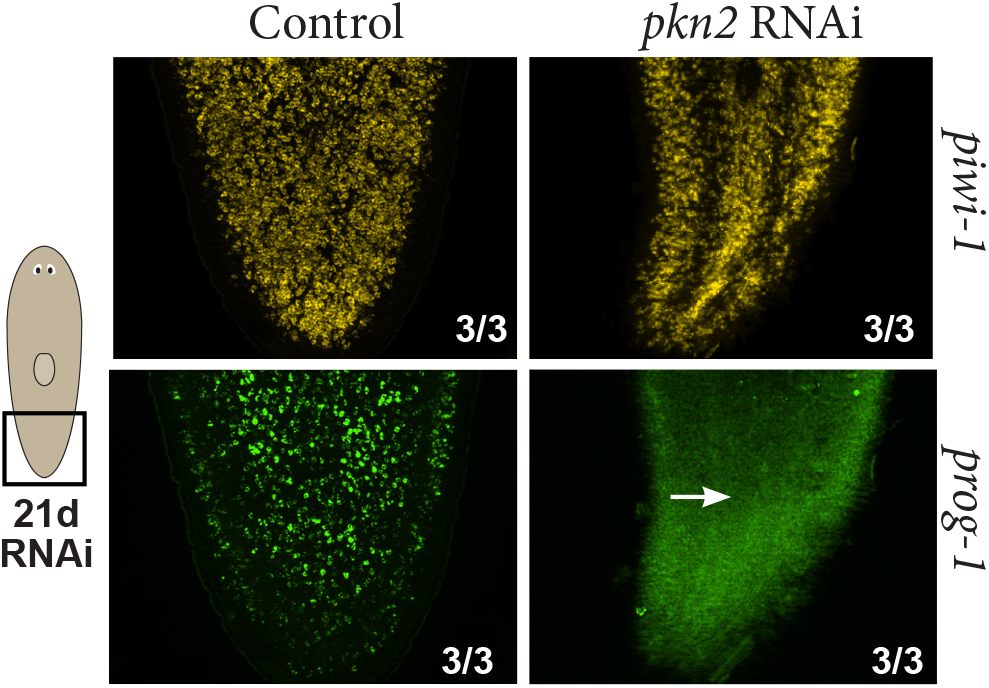
Left, animals were homeostatically administered either *pkn2* dsRNA or control dsRNA (6 dsRNA feedings over 21 days) then fixed and analyzed by FISH for expression of *piwi-1* and *prog-1. pkn2* RNAi caused loss of *prog-1* cells without reducing *piwi-1+* cells, suggesting this factor is required for epidermal differentiation.

